# Emergence of phenotypically distinct subpopulations is a factor in adaptation of recombinant *Saccharomyces cerevisiae* under glucose-limited conditions

**DOI:** 10.1101/2021.01.18.427050

**Authors:** Naia Risager Wright, Tune Wulff, Christopher T. Workman, Nanna Petersen Rønnest, Nikolaus Sonnenschein

**Author notes:** Correspondence: Nikolaus Sonnenschein, Technical University of Denmark.

## Abstract

Cells cultured in a nutrient-limited environment can undergo adaptation, which confers improved fitness under long-term energy limitation. We have previously shown how a recombinant *S. cerevisiae* strain, producing a heterologous insulin product, under glucose-limited conditions adapts over time at the average population level.

In this paper, we investigated this adaptation at the single-cell level by application of FACS and showed that three apparent phenotypes underlie the adaptive response observed at the bulk level: (1) cells that drastically reduced insulin production (23 %), (2) cells with reduced enzymatic capacity in central carbon metabolism (46 %), (3) cells that exhibited pseudohyphal growth (31 %). We speculate that the phenotypic heterogeneity is a result of different mechanisms to increase fitness. Cells with reduced insulin productivity have increased fitness by reducing the burden of the heterologous insulin production and the populations with reduced enzymatic capacity of the central carbon metabolism and pseudohyphal growth have increased fitness towards the glucose-limited conditions.

The results highlight the importance of considering population heterogeneity when studying adaptation and evolution.

## 1. Introduction

A growing number of pharmaceuticals, food ingredients and other valuable chemicals are today produced commercially by recombinant microbial cell factories. Expression of heterologous proteins confers a burden to the host strain which can result in an adverse selection towards low-producing cells during production (Rugbjerg & Olsson, 2020).

The chemostat mode of cultivation is an efficient strategy for industrial production processes, especially in continuous manufacturing, as the cells in the bioreactor can be kept in a constant growth environment for long periods with a constant output (Wright, Rønnest, & Sonnenschein, 2020). In this steady-state growth environment, the cells are exposed to nutrient limitation, e.g. glucose, either as a kinetic limitation of the cellular transport of nutrients or as limitation of the metabolic enzymes converting the nutrients (Schreiber & Ackermann, 2020). The nutrient-limited conditions in the chemostat impose a constant selective pressure on the production organism. Isogenic cells cultured in glucose-limited environments often undergo adaptation increasing fitness benefits under long-term energy limitation.

We have previously shown how a recombinant *S. cerevisiae* strain producing a heterologous protein adapts over time with respect to the population average protein levels (Wright et al., 2020). Over approximately 30 generations of glucose-limited growth, we observed a drastic decrease in recombinant protein production to almost half of the maximum value together with significant changes in the intracellular proteome. The underlying mechanisms of this adaptation are currently unknown but could not be associated to genetic mutations (unpublished results).

In nature, microorganisms live in complex communities consisting of many different species, subspecies and genetic variants. Phenotypic heterogeneity can occur in populations of genetically identical cells with respect to different traits including metabolism and morphology (Davis & Isberg, 2016). Phenotypic population heterogeneity with respect to metabolic activity has been shown in chemostat cultivations of microbes (Kundu, Weber, Griebler, & Elsner, 2020; Maharjan, Seeto, & Ferenci, 2007).

In this study, we have investigated the adaptation of a recombinant *S. cerevisiae* strain in glucose-limited cultures at the single-cell level. The results show that the bulk adaptive outcomes observed at the culture level, i.e., protein levels, physiology changes, and loss of productivity, are a mixed response composed of at least three apparent phenotypes or subpopulations. The results highlight the importance of considering population heterogeneity when studying adaptation.

## 2. Materials and Methods

### 2.1 Strains and experimental overview

An isogenic culture of the recombinant *S. cerevisiae* strain, C.U17 was cultured in prolonged chemostat cultivations (Seresht et al., 2013). The strain contains a 2μm vector with an expression cassette encoding the Triose Phosphate Isomerase 1 (TPI1) promoter and a gene encoding a single-chain insulin precursor. Both HIS3 and URA3 were used as auxotrophic selection markers. The C.U17 strain is referred to as the *initial cell clone* throughout the rest of the manuscript. After 271 hours of glucose-limited growth, end sample cells from one of the cultivations were collected and stored as glycerol stocks at −80°C. These cells are referred to as *end sample cells*. One of the glycerol stocks were thawed, washed three times in PBS buffer and used for FACS sorting (Fluorescence Activated Cell Sorting) of three individual populations separated based on particle size measured as forward scatter light area (FSC-A) (Figure 1A). 10,000 sorted cells from each population were propagated in individual shake flasks with minimal medium at 30°C, harvested at an OD_600_ of 20 and stored as glycerol stocks at −80°C (Figure 1B). The glycerol stocks were used to reinitiate new chemostat cultures with each of the FACS sorted populations (Figure 1C).

**Figure 1:**
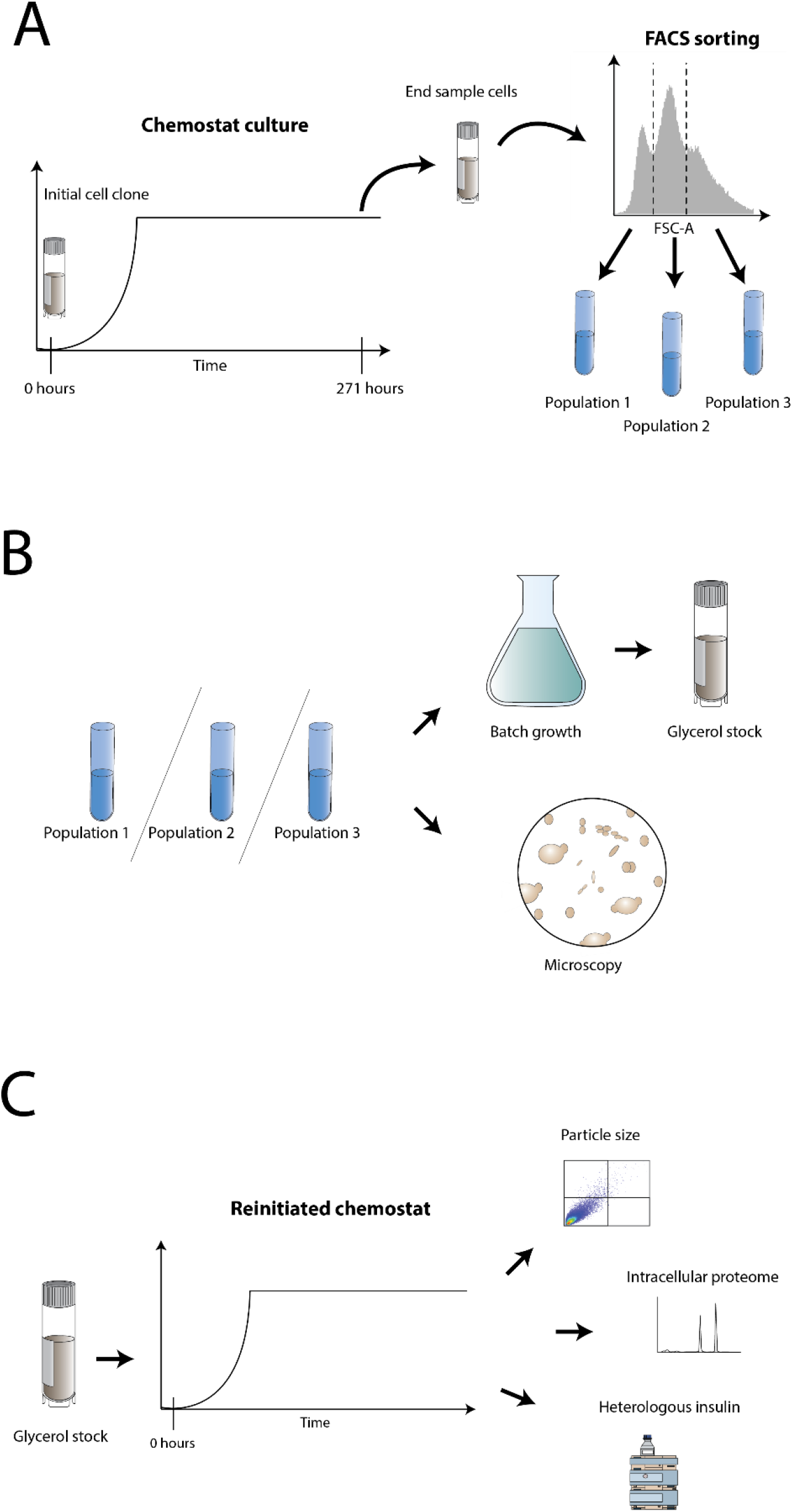
Overview of the experimental setup. A) The monoclonal initial cell clone was used to inoculate a chemostat culture. After 271 hours of continuous cultivation a sample of adapted cells were taken (end sample cells) and stored as glycerol stock. The end sample cells were further separated by FACS sorting into three different subpopulations based on FSC-A. B) The FACS sorted subpopulations were inspected by microscopy, propagated in batch cultures and stored as glycerol stock. C) Each of the FACS sorted subpopulations were cultured in reinitiated chemostats and compared in terms of particle size (FSC-A), intracellular proteome and heterologous insulin production. In addition, the end sample cells were cultured in reinitiated chemostats and analyzed for heterologous insulin production and particle size (FSC-A).

### 2.2 Chemostat cultivations

Aerobic chemostat cultivations were performed in 0.5 L fully instrumented and automatically controlled BIOSTAT^®^ reactors (Sartorius Stedim Biotech S.A, Germany). The strains were cultured in duplicates as previously described in Wright et al., (2020) at a temperature of 28 °C, pH of 5.9, aeration rate of 2 vvm and dilution rate of 0.1 h^−1^. A minimal medium with a glucose concentration of 75 g/L was used. Each cultivation was initiated with 8 hours batch phase and 52 hours fed-batch phase. Cell dry weight and extracellular heterologous insulin production were measured each day as previously described in Wright et al., (2020).

### 2.3 Fluorescent activated cell sorting (FACS)

Samples for flow cytometry analysis were collected every day and stored in glycerol at −80 °C. Prior to analysis, the samples were thawed and washed three times in PBS buffer. Flow cytometry analysis and cell sorting with respect to particle size (FSC-A) were performed using a Sony Cell Sorter SH800S. 100.000 cells were analyzed in each sample using a 100 μm microfluidic sorting chip. The samples were diluted to obtain an event rate below 1000 eps. The raw flow cytometry data (fcs files) were analyzed in the software environment R version 3.6.1 using the flowCore package (Ellis et al., 2020). Stacked density plots of the log2(FSC-A) distribution at different time points were constructed by application of the ggplot2 package in R (Wickham, 2016). Histogram bar charts of log2(FSC-A) were constructed by sorting the cell count data into 833 uniformly sized bins using the build-in *hist* function in R.

### 2.4 Analysis of intracellular proteins

A minimum of four samples were withdrawn for analysis of intracellular proteins at different time points of the reinitiated chemostat cultivations of the three FACS sorted populations (See Supplementary materials Table S1 for an overview of the different samples). The samples were stored at −80 °C before further processing. Intracellular proteins were quantified by label-free quantification as previously described in Wright et al., (2020). For analysis of the samples, liquid chromatography was performed on a CapLC system (Thermo Fisher Scientific) coupled to an Exploris 480 mass spectrometer (Thermo Fisher Scientific). The peptides were separated with a flow rate of 1.2 μl/min on a 75-μm × 15 cm 2 μm C18 easy spray column. A stepped gradient, going from 4% to 40% acetonitrile in water over 50 minutes was applied. Mass spectrometry (MS)-level scans were performed with the following settings: Orbitrap resolution: 60,000; AGC Target: 1.0e6; maximum injection time: 50 ms; intensity threshold: 5.0e3; and dynamic exclusion: 25 s. Data-dependent MS2 selection was performed in Top 12 mode with HCD collision energy set to 30 % (AGC target: 1.0e4; maximum injection time: 22 ms).

### 2.5 Data processing of proteome data

For analysis of the thermos rawfiles, Proteome Discover 2.3 (Thermo Fisher Scientific) was applied. The following settings were used for the analysis: Fixed modifications: Carbamidomethyl (C) and Variable modifications: oxidation of methionine residues. First search mass tolerance of 20 ppm and an MS/MS tolerance of 20 ppm. Trypsin was selected as an enzyme and allowing one missed cleavage. False discovery rate was set at 0.1%. The data was searched against the *S. cerevisiae* database retrieved from Uniprot with proteome ID AUP000002311 and the sequence of the heterologous insulin. The data sets can be found at data.dtu.dk (https://doi.org/10.11583/DTU.13536179).

Batch variations between different proteome datasets were reduced by scaling each protein such that the mean log2(abundance) was the same between data sets. A differential expression analysis was performed between *Population 1, Population 2* and *Population 3* for samples taken in the beginning of the cultures (≤ 48 hours of chemostat growth) and again in the end of the cultures (after 254 hours of chemostat growth). Only proteins which were measured in all samples between the compared populations were included in the analysis meaning that 2635 proteins were compared between *Population 1* and *Population 3*, 2811 proteins were compared between *Population 1* and *Population 2* and 2679 proteins were compared between *Population 2* and *Population 3*. The analysis was performed using the EdgeR package (Robinson, McCarthy, & Smyth, 2010) in R version 3.6.1. The proteome from the beginning of chemostat cultures with the three populations were furthermore compared to a previously published proteome of the *initial cell clone* (Wright et al., 2020). For an overview of the samples used for the comparison, see Supplementary materials Table S2. 2716 proteins between *Population 1* and the *initial cell clone* were compared, 2770 proteins between *Population 2* and the *initial cell clone* were compared and 2692 proteins were included in the comparison of *Population 3* and the *initial cell clone*.

For each comparison between two *strains*, proteins were grouped in clusters depending on whether the level of the proteins were higher (log2 fold-change > 0.5, q-value < 0.05) or lower (log2 fold-change < 0.5, q-value < 0.05) in strain A compared to strain B. Gene ontology (GO) process terms were obtained from geneontology.org/annotations/sgd.gaf.gz on 16 November, 2020. A one-sided Fisher’s exact test was used to investigate whether the protein clusters, were enriched with proteins annotated with certain GO process terms (q-value <0.05). The test was performed using the R package *bc3net* package (de Matos Simoes, Tripathi, & Emmert-Streib, 2012).

### 2.6 Microscopy

The morphology of FACS sorted populations were visually inspected using a LMI-005-Leica Microscope and a Confocal Microscope-SP8.

### 2.7 Determination of maximum growth rate in batch cultures

Maximum growth rates were determined based on OD_600_ measurements from exponentially growing cells in 100 ml shake flasks with minimal medium and 3 % v/v glucose (Seresht et al., 2013). Three biological replicates were performed for each strain.

## 3 Results & Discussion

### 3.1 FACS analysis of chemostat cultivations with *S. cerevisiae,* C.U17

We cultivated the recombinant *S. cerevisiae* strain, C.U17, in duplicated chemostats and performed time-resolved analysis of the population with respect to particle size measured by FSC-A (used as a proxy for Morphology), (Figure 1A). We observed the differentiation of three main subpopulations over time with respect to FSC-A (Figure 2; See Supplementary materials Figure S1 and S6 for replicates). We named the three subpopulations *Population 1* (small particles), *Population 2* (medium particles) and *Population 3* (large particles), respectively. *Population 1* corresponded to 23 % of the total population after 270 hours of glucose-limited growth, *Population 2* corresponded to 46 % and *Population 3* corresponded to 31 % (Figure 3A).

**Figure 2:**
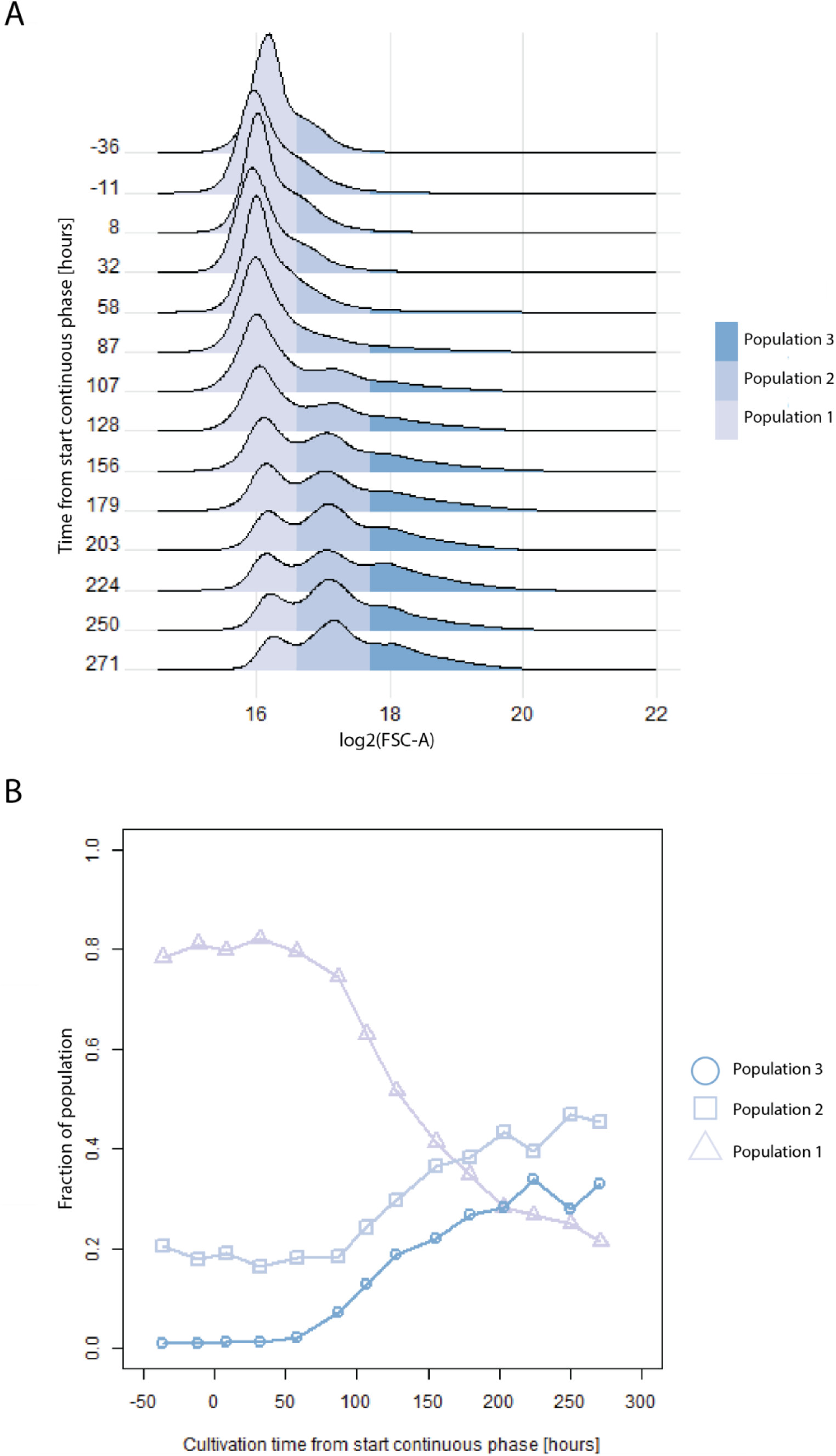
Distribution of cells with FSC-A corresponding to Population 1 (log2(FSC-A) < 16.6), Population 2 (16.6 < log2(FSC-A) < 17.7) or Population 3 (log2(FSC-A) > 17.7) at different time points during a representative culture of the initial cell clone. A) Density plot of log2(FSC-A) at different cultivation time points. 100,000 cells were analyzed at each time point. B) Fraction of cells with a FSC-A corresponding to each of the three populations as function of cultivation time.

**Figure 3:**
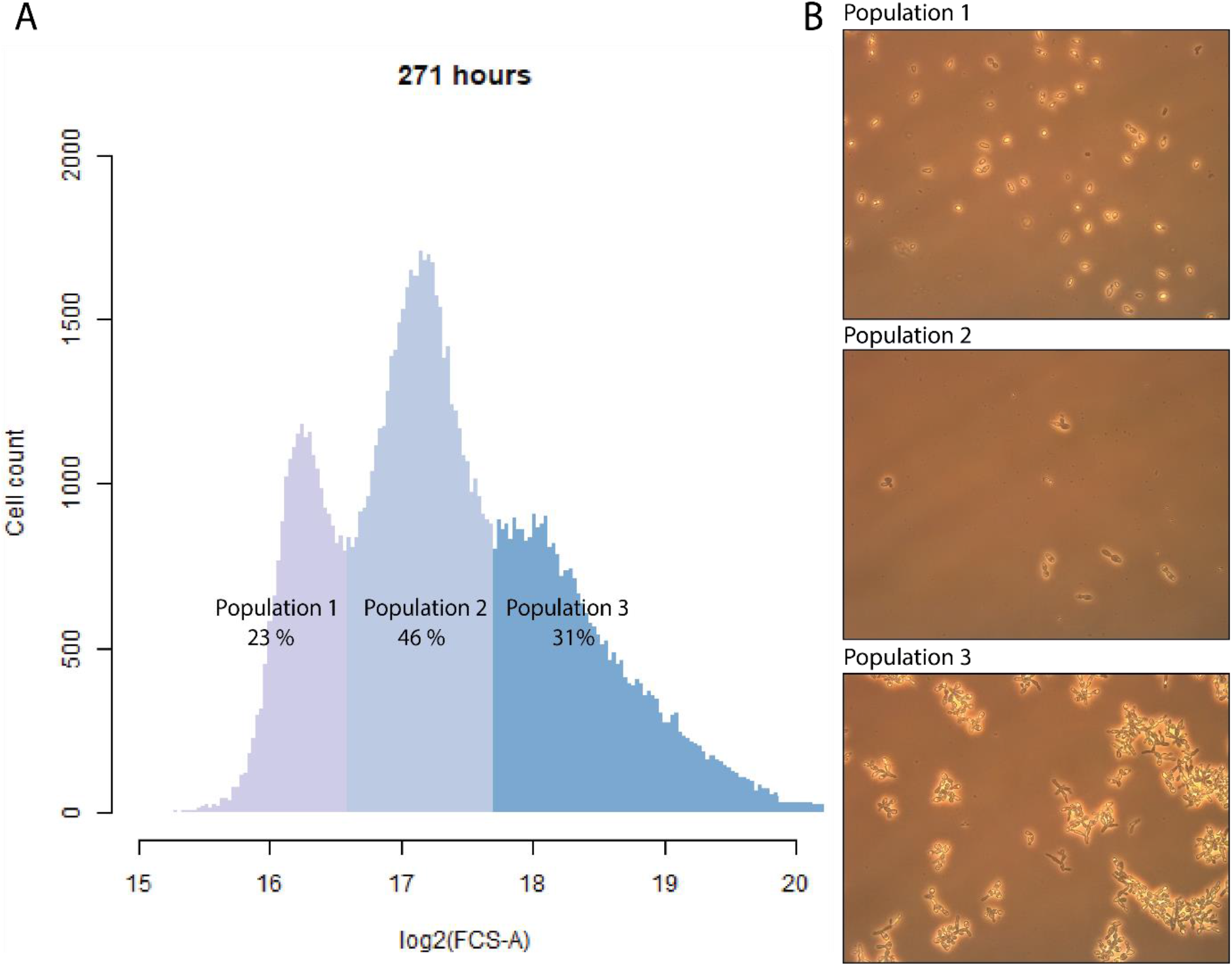
A) Cell count with respect to log2(FCS-A) measured by FACS after 271 hours of chemostat culture with the initial cell clone. The color code indicates the separation of cells into three subpopulations for FACS sorting. 100,000 cells were analyzed. B) Images of cell morphology of the three FACS sorted subpopulations.

### 3.2 Morphology of FACS sorted populations

We FACS sorted each of the three subpopulations based on FSC-A and inspected the subpopulations by microscopy (Figure 1A, B). We observed that *Population 1* primarily contained single cells where about a fourth presented small round buds, whereas *Population 2* consisted of multi-budded cells with a more ellipsoidal cell shape (Figure 3B). *Population 3* contained branches of elongated cells with multiple buds (pseudohyphae).

Morphological changes towards a more filamentous and pseudohyphal growth are known effects of chemostat growth and also a known adaptive response of cells in a nutrient poor environment as a strategy to forage for nutrients (Ceccato-Antonini and Sudbery 2004; Hope et al. 2017; Rai et al. 2019). This suggests that *Population 3* has emerged as a result of the glucose-limited conditions.

Flocculation and wall growth have been observed in prolonged chemostats where cells stick to the surface of the culture vessels (Dykhuizen & Hartl, 1983). However, wall growth has not been observed in this study.

Morphological changes exhibited by cell subpopulations comprising large single-budded and multi-budded cells have previously been observed in industrial scale chemostat cultivations with *S. cerevisiae* and were related to hypoxia (Aon, Tecson, & Loladze, 2018). In the current study, no sign of oxygen limitation can be found during the chemostat cultures (Figure 4). Thus, we link the formation of the subpopulations to the selective pressure of the constant glucose-limited environment. We confirm this link by glucose-pulse experiments. A uniform particle size and morphology with no significant changes over time could be detected in cultures exposed to continuous glucose pulsing (Supplementary materials Figure S11, S12, S13).

**Figure 4:**
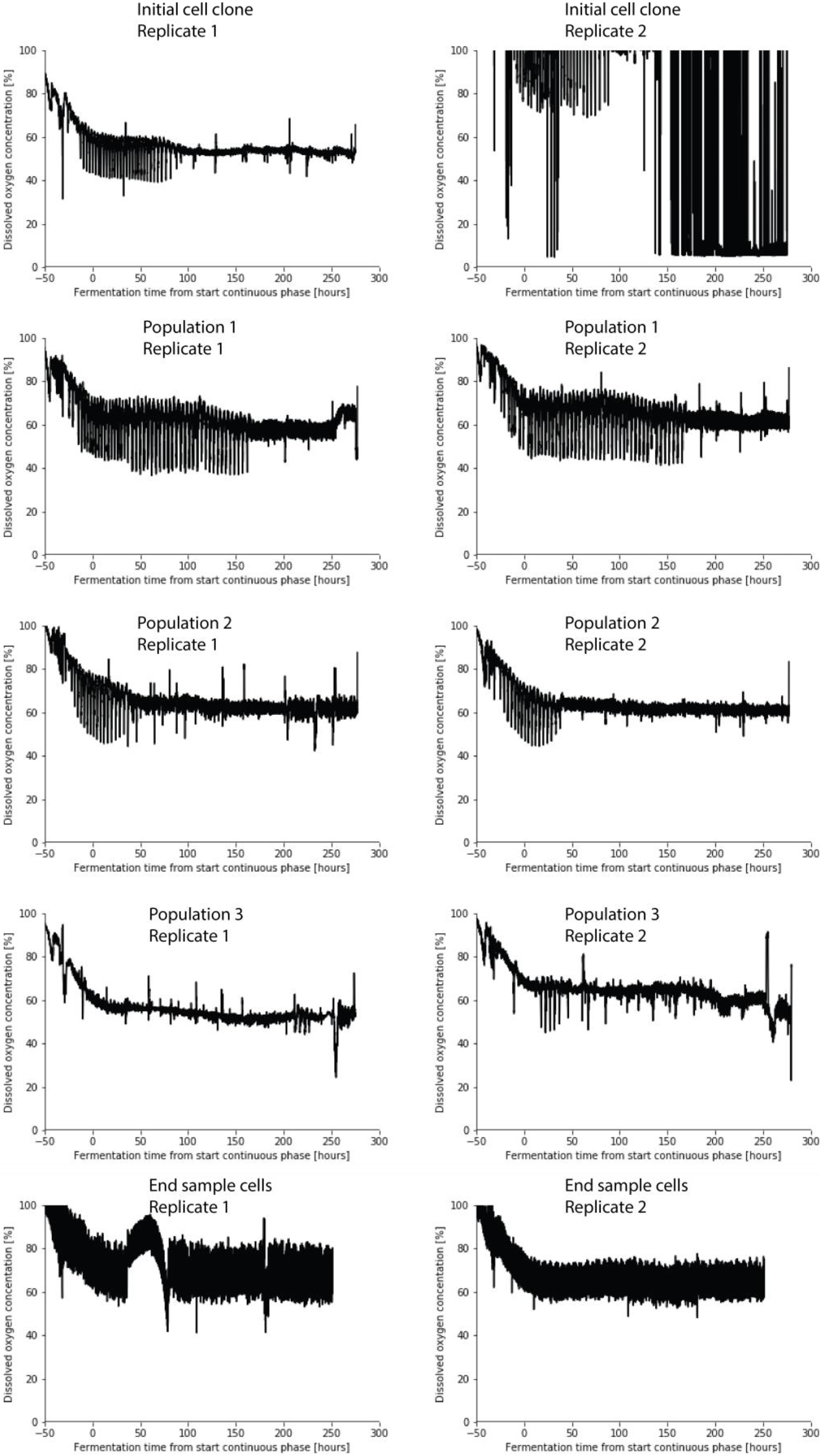
Dissolved oxygen tension during chemostat cultivations with the Initial cells clone, the end sample cells and the sorted subpopulations population1, Population 2 and Population 3. The pO_2_ electrode was defective for replicate 2 with the initial cell clone.

### 3.3 Maximum growth rate of FACS sorted subpopulations

Each of the FACS sorted individual populations had maximum growth rates equal to or higher than the *initial cell clone* (Figure 5A). *Population 1* had a significant higher maximum growth rate on glucose compared to the other populations, which can be explained by a lower metabolic burden of the heterologous insulin production (this is further discussed in section 3.4.2).

**Figure 5:**
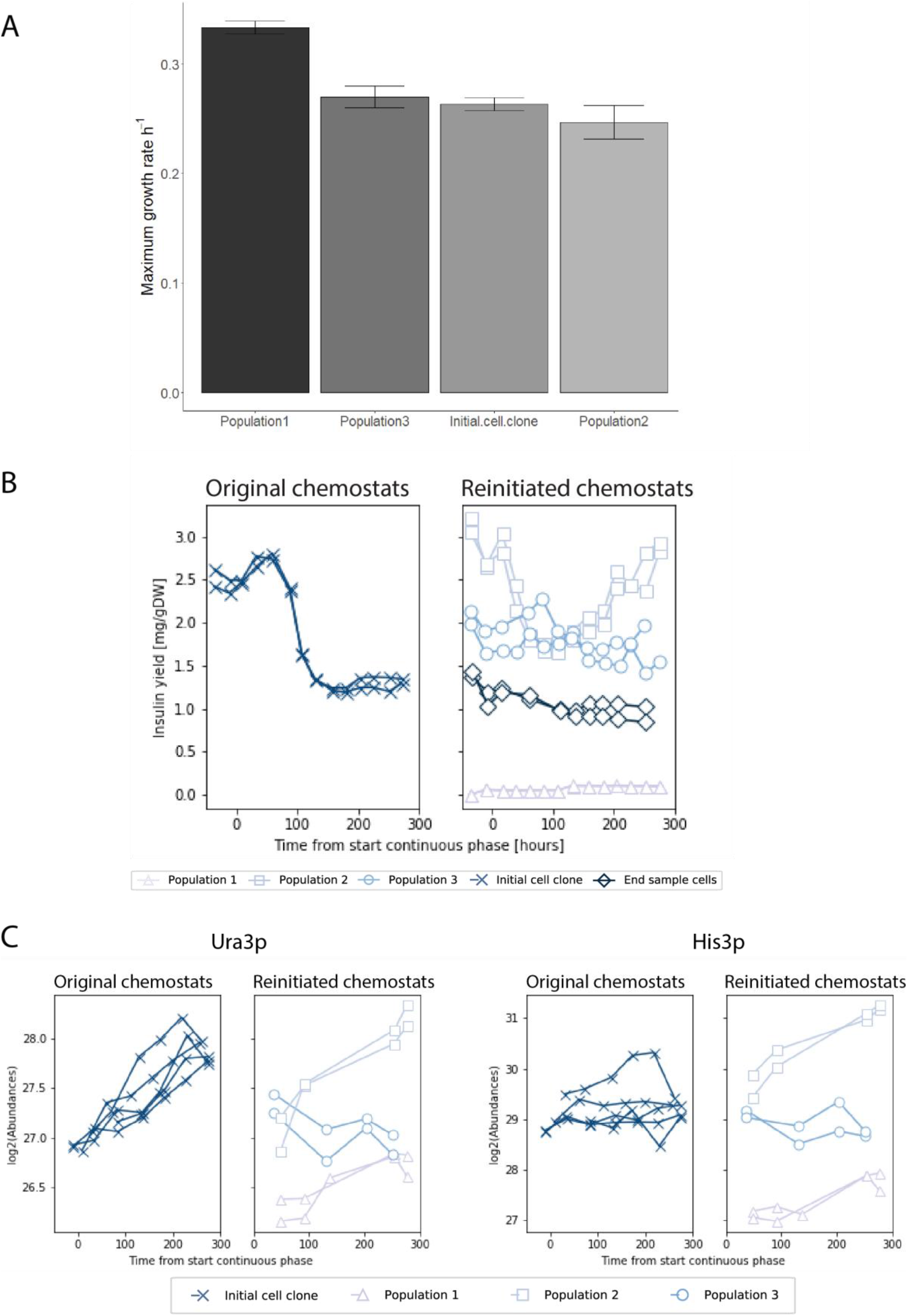
A) Maximum growth rate in batch cultures. Error bars indicate differences between three biological replicates, B) Insulin yield over time in chemostat cultivations. C) Levels of His3p and Ura3p over time in chemostat cultivations.

### 3.4 Reinitiated chemostat cultures of FACS subpopulations

Each of the FACS sorted subpopulations were propagated in batch cultures and stored as glycerol stocks (Figure 1A, B).

We cultured each of the propagated populations in new chemostat cultures in order to characterize the populations individually, with respect to particle size (FSC-A), heterologous insulin production and intracellular proteome (Figure 1C). We also compared the subpopulations to the *end sample cells* from which they were sorted with respect to heterologous insulin production. Biomass concentrations measured as cell dry weight for each cultivation can be found in Supplementary materials Figure S17.

#### 3.4.1 Particle size (FSC-A)

We observed that the particle size of some of the cells in *Population 2* and *Population 3* after propagation in batch cultures had changed towards particle sizes typical of *Population 1* (Supplementary materials Figure S2, S3, S4, S7, S8, S9). However, the particle size increased again over time in the reinitiated chemostats.

#### 3.4.2 Heterologous insulin production

The production of the recombinant insulin decreased over time when the *initial cell clone* was cultivated under prolonged glucose-limited conditions (Figure 5B). We have previously coupled this decline to changes in the intracellular proteome of the strain observed in measurements at the culture level (Wright et al., 2020). The *end sample cells*, continued to produce insulin at the same low level when cultured *de novo* in chemostats, while the subpopulations showed divergent phenotypes with respect to heterologous insulin production (Figure 5B). In the beginning of the reinitiated cultures, the sum of the insulin yields of *Population 1-3* was ~1.8 mg/gDW (adjusting for the ratios between the three subpopulations at the end of the cultivations with the *Initial cell clone*). This yield approximates the insulin yield observed by bulk measurements in the end of the cultivations with the *initial cell clone*. This indicates that the insulin yield of the three populations were conserved during the batch propagation of the FACS sorted cells (estimated to more than 40 generation).

Only small amounts of insulin could be detected in cultures with *Population 1*. The production of a heterologous protein is a burden for the cells (Peebo & Neubauer, 2018) and cells with reduced productivity will have a growth advantage compared to cells, which are not able to adapt (Rugbjerg & Olsson, 2020). We observe this advantage as a higher maximum growth rate for *Population 1* under glucose rich conditions (Figure 5A). Therefore, we suggest that *Population 1* has mainly arisen due to the burden of the heterologous protein production and to a lesser extent due to the glucose-limited conditions. The determined maximum growth rate for *Population 1* is slightly lower than the growth rate of the wildtype CEN.PK113-7D strain (0.37 h^−1^ ± 0.01), which has previously been used as reference strain for comparison with the C.U17 strain (Seresht et al., 2013).

The bulk measurements of insulin in the *initial cell clone* reached a new steady state after around 150 hours of glucose-limited growth (Figure 5B). Moreover, a steady state between the three morphological phenotypes seemed to occur after around 200 hours of chemostat cultivation (Figure 2B). Cultivations with the *end sample cells* for another 250 hours under glucose-limited conditions confirm the steady state with respect to insulin production (Figure 5B). This indicates that none of the three populations had a significant higher fitness under chemostat conditions compared to the other. Based on the maximum growth rate experiments, we would have expected that *Population 1* should take over the entire population in the chemostat and that *Population 2* and *Population 3* gradually would be washed out. However, the increase in fitness of chemostat-adapted cells is often specific to the low nutrient-limited environment and many organisms adapted to chemostat conditions show reduced growth capabilities in nutrient rich environments (Gresham & Hong, 2015; Hong & Gresham, 2014; Wenger et al., 2011). Thus, a higher maximum growth rate under glucose rich conditions will not be an adequate measure of fitness under glucose-limited conditions.

Yeast cells release a variety of different metabolites and prefer the uptake of extracellular metabolites over self-synthesis (Campbell et al., 2015). Population heterogeneity can emerge as a consequence of metabolic cooperation between cells and a population as a whole can benefit from division of labor between individuals (Ackermann, 2015; Campbell, Vowinckel, & Ralser, 2016). We speculate that the three populations in the chemostat cooperate in metabolism and exchange metabolites and that this may explain the observed steady state between the subpopulations. However, this suggestion needs to be further investigated.

#### 3.4.3 Intracellular proteome

The FACS sorted cells were further characterized with respect to their intracellular proteome during chemostat growth. We compared the intracellular proteome in the beginning of chemostat cultures of the three populations with our previously reported proteomics dataset from the beginning of chemostat cultures with the *initial cell clone* (Wright et al., 2020), (Table 1). Moreover, we compared the proteome of the three individual populations in the beginning of the chemostat cultures and again in the end of the cultivations.

**Table 1:** Comparison of the proteome during chemostat growth of the initial cell clone and Population 1-3. The table illustrates the fraction of differentially expressed proteins for each comparison (log2 fold change > 0.5 or log2 fold change < −0.5, q-value < 0.05). Moreover, GO process terms, which based on a Fischer’s exact test, are enriched by proteins in a cluster of proteins with q-values <0.05 and log2 fold-change >0.5 are shown. In a comparison of two strains A vs. B, the protein levels of A is higher than the protein levels of B if log2 fold-change > 0. No differentially expressed GO-terms could be found for clusters of proteins with log2 fold-change <0.5 and q-values < 0.05 for any of the comparisons.

| Comparison | Fraction of measured proteome which is differentially expressed ($q$-value $< 0.05$, $\log_2$ fold-change $> 0.5$ or $\log_2$ fold change $< -0.5$) | Enriched GO process terms in a cluster of proteins with $\log_2$ fold-change $> 0.5$, $q$-value $< 0.05$ ($q$-value $< 0.05$) |
| --- | --- | --- |
| Initial cell clone vs Population 1<br>Beginning of cultivation ($\leq 48$ hours of chemostat growth) | 6 % | - |
| Initial cell clone vs Population 2<br>Beginning of cultivation ($\leq 48$ hours of chemostat growth) | 13 % | Carbohydrate metabolic process, Oxidation-reduction process, Gluconeogenesis, Glycolytic process, NADH oxidation |
| Initial cell clone vs Population 3<br>Beginning of cultivation ($\leq 48$ hours of chemostat growth) | 4 % | Oxidation-reduction process |
| Population 1 vs Population 2<br>Beginning of cultivation ($\leq 48$ hours of chemostat growth) | 14 % | Oxidation-reduction process, Tricarboxylic acid cycle, Fatty acid metabolic process, Fatty acid beta-oxidation, Gluconeogenesis, Glyoxylate cycle, NADH oxidation, lipid metabolic process, carboxylic acid metabolic process, amino acid catabolic process to alcohol via Ehrlich pathway, malate metabolic process, isocitrate metabolic process, cellular respiration |
| Population 1 vs Population 3<br>Beginning of cultivation ($\leq 48$ hours of chemostat growth) | 2 % | - |
| Population 2 vs Population 3<br>Beginning of cultivation ($\leq 48$ hours of chemostat growth) | 0 % | - |
| Population 1 vs Population 2<br>End of cultivation (254 hours of chemostat growth) | 16 % | - |
| Population 1 vs Population 3<br>End of cultivation (254 hours of chemostat growth) | 0.6 % | - |
| Population 2 vs<br>Population 3<br>End of cultivation<br>(254 hours of<br>chemostat growth) | 0.07 % | - |

In the beginning of the chemostat cultures, *Population 2* differed from *Population 1* and the *initial cell clone* with respect to proteins involved in the central carbon metabolism (Table 1-3). The levels of these proteins were significantly lower in *Population 2* (See Figure 6 for example of glycolytic proteins). Protein synthesis is an energetically expensive process. Cells, which can economize the protein synthesis, e.g., by reducing the production of overexpressed proteins in a nutrient-limited environment, will have an advantage over cells, which cannot adapt. Reduced capacity of the central carbon metabolism including the glycolysis and TCA cycle is a well-known adaptive response to chemostat conditions (Franchini & Egli, 2006; Jansen et al., 2005; Mashego, Jansen, Vinke, van Gulik, & Heijnen, 2005). This suggests that the establishment of *Population 2* is a response to the glucose-limited conditions and confers a fitness advantage in the chemostat.

**Table 2:** GO process terms, which based on a Fischer’s exact test are enriched by proteins in a cluster of proteins with significant lower levels in Population 2 compared to the initial cell clone in the beginning of cultivations (≤ 48 hours of chemostat growth), (log2 fold-change > 0.5, q-value < 0.05). Proteins from the cluster are linked to the specific GO process terms, which they belong to.

| GO description | q-value | Proteins |
| --- | --- | --- |
| Carbohydrate metabolic process | 4.73E-06 | Ath1p, Cts1p, Exg1p, Fba1p, Glc3p, Glk1p, Gpd1p, Gph1p, Gut1p, Ima5p, Mdh1p, Mdh2p, Nqm1p, Pgm2p, Snf4p, Sol1p, Sol4p, Suc2p, Ydr248cp |
| Oxidation-reduction process | 1.08E-04 | Adh1p, Adh2p, Adh5p, Ahp1, Ald4p, Arg5,6p, Ayr1p, Bdh1p, Bna1p, Cta1p, Cyb2p, Ecm4p, Erg11p, Fas1p, Fas2p, Fdh1p, Fox2p, Gcv2p, Gcy1p, Gnd2p, Gpd1p, Gpx2p, Gre2p, Gut2p, His4p, Hom2p, Idp2p, Idp3p, Mdh1p, Mdh2p, Nde2p, Oye3p, Pgk1p, Pox1p, Prx1p, Qcr6p, Rnr2p, Rnr4p, Tdh1p, Ydl124wp, Yjr096wp, Yml131wp |
| Gluconeogenesis | 5.62E-04 | Eno1p, Eno2p, Fba1p, Gpm1p, Gpm2p, Mdh2p, Pgk1p, Tdh1p, Tpi1p |
| Glycolytic process | 7.71E-03 | Eno1p, Eno2p, Fba1p, Glk1p, Gpm1p, Gpm2p, Pgk1p, Tdh1p, Tpi1p |
| NADH oxidation | 1.88E-02 | Adh1p, Adh2p, Adh5p, Gpd1p, Gut2p, Nde2p |

**Table 3:** GO process terms, which based on a Fischer’s exact test are enriched by proteins in a cluster of proteins with significant lower levels in Population 2 compared to Population 1 in the beginning of cultivations (≤ 48 hours of chemostat growth), (log2 fold-change > 0.5, q-value < 0.05)). The proteins from the cluster are linked to the specific GO process terms, which they belong to.

| GO description | q-value | Proteins |
| --- | --- | --- |
| **Oxidation-reduction process** | 6.55E-06 | Fas2p, Fas1p, Ald4p, Pgk1p, Tdh2p, Fdh1p, Fox2p, Tdh1p, Kgd1p, Gut2p, Cyb2p, Idp2p, Lys9p, Cta1p, Arg5,6p, His4p, Yhb1p, Adh2p, Adh1p, Mdh1p, Mcr1p, Pox1p, Idh1p, Ndi1p, Ydl124wp, Hom2p, Mdh3p, Sod1p, Gdh2p, Gcy1p, Mae1p, Sdh1p, Dld1p, Prx1p, Idp3p, Gnd2p, Rnr4p, Idh2p, Adh3p, Bna1p, Sdh2p, Erg11p, Gcv1p, Ahp1p, Cir2p, Gpd1p, Cyt1p, Zta1p, Rip1p, Yml131wp, Rnr1p, Mdh2p, Gre2p, Nde2p, Adh5p, Bdh1p, Bna4p, Sdh4p, Sdh3p, Ifa38p, Met12p |
| **Tricarboxylic acid cycle** | 9.33E-06 | Aco1p, Kgd1p, Idp2p, Cit1p, Mdh1p, Idh1p, Lsc2p, Mdh3p, Sdh1p, Idp3p, Idh2p, Mls1p, Cit2p, Sdh2p, Icl1p, Cit3p, Mdh2p, Sdh4p, Sdh3p |
| **Fatty acid metabolic process** | 1.63E-05 | Fas2p, Fas1p, Fox2p, Pox1p, Pot1p, Faa2p, Cat2p, Faa4p, Yat2p, Yat1p, Faa3p, Dci1p, Ifa38p |
| **Fatty acid beta-oxidation** | 8.48E-04 | Fox2p, Pox1p, Mdh3p, Pot1p, Idp3p, Tes1p, Eci1p |
| **Gluconeogenesis** | 1.24E-03 | Pgk1p, Eno2p, Tdh2p, Fba1p, Eno1p, Tdh1p, Gpm1p, Tpi1p, Pyc2p, Mdh2p |
| **Glyoxylate cycle** | 2.31E-03 | Idp2p, Mdh3p, Idp3p, Icl2p, Mls1p, Cit2p, Icl1p |
| **Glycolytic process** | 5.14E-03 | Pgk1p, Eno2p, Cdc19p, Tdh2p, Fba1p, Eno1p, Glk1p, Tdh1p, Gpm1p, Tpi1p |
| **NADH oxidation** | 1.15E-02 | Gut2p, Adh2p, Adh1p, Ndi1p, Adh3p, Gpd1p, Nde2p, Adh5p |
| **Lipid metabolic process** | 1.11E-02 | Fas2p, Fas1p, Fox2p, Ino1p, Pox1p, Pot1p, Faa2p, Gpt2p, Cat2p, Faa4p, Erg11p, Yat2p, Yat1p, Faa3p, Dci1p, Plb2p, Ifa38p, Tgl1p |
| **Carboxylic acid metabolic process** | 2.98E-02 | Mdh1p, Mdh3p, Icl2p, Leu9p, Icl1p, Mdh2p |
| **Amino acid catabolic process to alcohol via Ehrlich pathway** | 4.69E-02 | Adh2p, Adh1p, Adh3p, Adh5p |
| **Malate metabolic process** | 4.69E-02 | Mdh1p, Mdh3p, Mae1p, Mdh2p |
| **Isocitrate metabolic process** | 4.69E-02 | Idp2p, Idh1p, Idp3p, Idh2p |
| **Cellular respiration** | 4.69E-02 | Sdh1p, Sdh2p, Sdh4p, Sdh3p |

**Figure 6:**
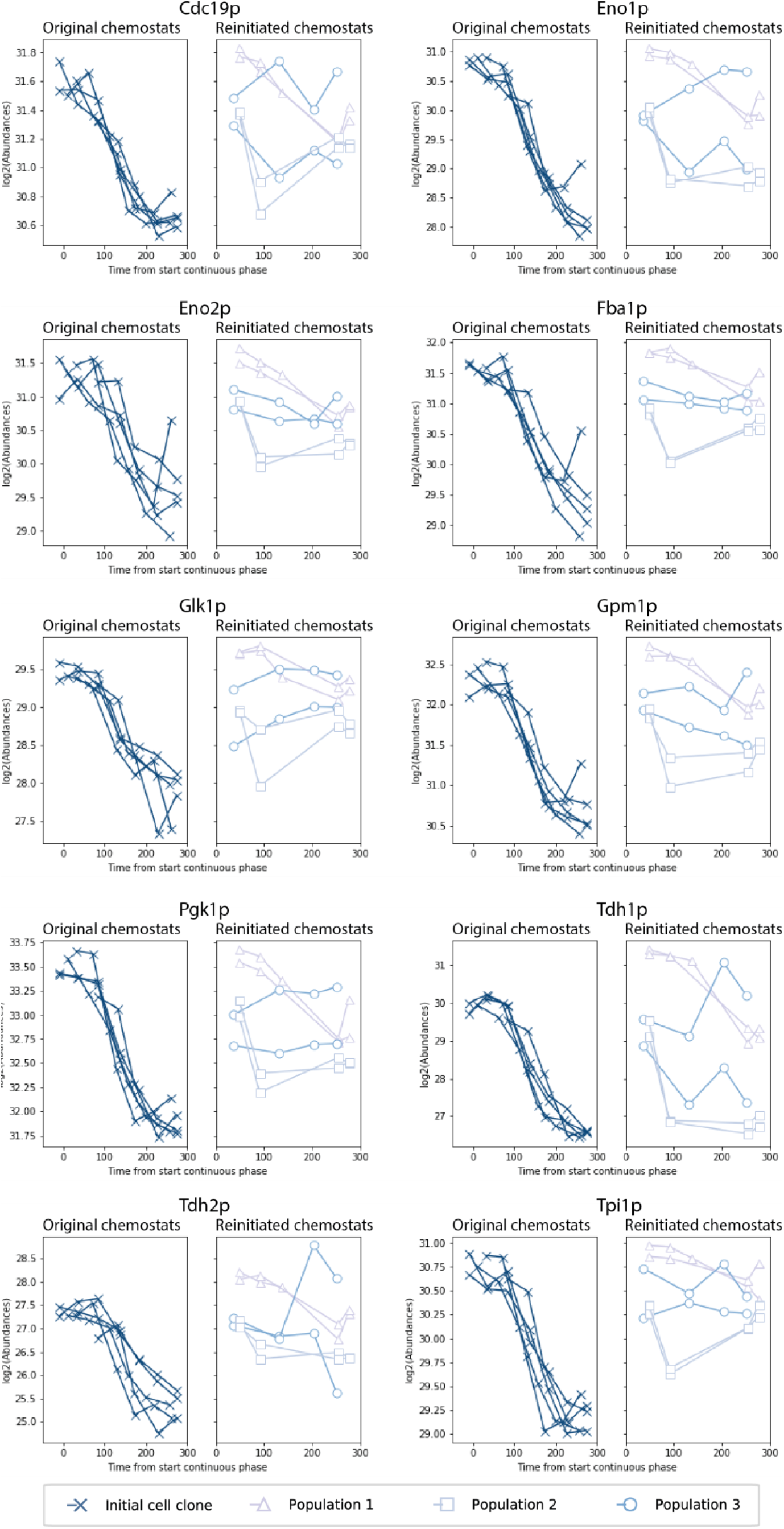
Levels of glycolytic enzymes measured during replicated chemostat cultivations with the initial cell clone, Population 1, Population 2 and Population 3. Only proteins, which differ significantly between Population 1 and Population 2 in the beginning of the cultivations (≤ 48 hours of chemostat growth) are pictured (q-value <0.05, log2 fold-change >0.5).

6 % of the measured proteome differed between *Population* 1 and the *initial cell clone* (log2 fold-change > 0.5 or log2 fold-change < −0.5, q-value <0.05), (Table 1). No differentially expressed GO terms could be found related to these proteins (q-value <0.05). However, levels of the proteins expressed from the selective markers, URA3 and HIS3 were significantly lower in cultivations with *Population 1* compared to cultivations with the *initial cell clone* and *Population 2* (Figure 5C). This indicates that either a reduced plasmid copy number or a down regulation of the plasmid genes caused the low insulin production in *Population 1*. A decline in plasmid copy number over cultivation time has previously been measured in the bulk of chemostat cultivations with *the initial cell clone* in the time frame investigated in this study (Seresht et al., 2013) and may be related to *Population 1* growing over time. In general, several proteins involved in the biosynthesis of uridine are significantly decreased in the beginning of cultivations with *Population 1* compared to *Population 2* (q-value<0.05) (see Supplementary materials Figure S16 for levels of the specific proteins). This is not the case for the biosynthesis pathway of histidine (see Supplementary materials Figure S15 for levels of the specific proteins).

After 271 hours of glucose-limited growth, only 16 proteins differed with more than 0.5 log2 fold-change between *Population 1* and *Population2* (q-value < 0.05), (Table 1, see Supplementary materials Table S3 for the specific proteins). This indicates that two adaptive mechanisms occurred over time in *Population 1*. In the original chemostat cultures, the fitness of the cells was increased towards the burden of the heterologous insulin production by a decrease of the insulin productivity. In the reinitiated chemostat cultures, the fitness was increased towards the glucose-limited conditions by a decrease in the overcapacity of especially enzymes linked to the central carbon metabolism. We have previously shown that the adaptation to glucose-limited conditions is enhanced by the production of a heterologous product (Wright et al., 2020). This may explain the delayed adaptation in *Population* 1 with respect to the reduction of enzymatic overcapacity compared to *Population 2*. During the first 100 hours of the reinitiated chemostats with *Population 2* there seems to be a further reduction in the glycolytic capacity of the cells (Figure 6).

No significantly expressed proteins could be found between *Population 2* and *Population 3* in the beginning of chemostat cultivations, whereas 50 proteins differed between *Population 1* and *Population 3* (log2 fold-change > 0.5 or log2 fold-change < −0.5, q-value <0.05). *Population 3* consisted of cells with multiple-buds that span a larger variety of cells with respect to particle size. Therefore, a larger variation was observed between replicates with this population (Supplementary materials Figure S14). This variation explains the lower level of significantly changing proteins between *Population 3* and the other subpopulations.

#### 3.4.4 Cell synchrony

We have previously shown that the *initial cell clone* synchronized growth during the first 100 hours of chemostat growth (Wright et al., 2020). The time point where the synchronized growth stopped correlated with the time point where the heterologous insulin production started to decrease (Figure 4; Figure 5B). This also correlated well with the time point where the population heterogeneity with respect to particle size (FSC-A) arose in the culture (Figure 2B). Neither the e*nd sample cells,* nor *Population 3* showed synchronized growth in the reinitiated cultures. We explain this by the larger fraction of population heterogeneity in the reinitiated chemostats already from the beginning of the cultures (Supplementary materials Figure S7-S10). This heterogeneity was conserved during the initial batch propagation. *Population 1* showed synchronized growth for 150 hours, whereas *Population 2* showed synchronized growth for 50 hours. The stop of synchronized growth of *Population 2* correlated with an increase in population heterogeneity in the reinitiated cultures with respect to particle size (FSC-A), (Supplementary materials Figure S8).

### 3.5 Three apparent phenotypes underlie the adaptive response observed at the bulk level

We have previously shown that the recombinant *S. cerevisiae* strain (C.U17) adapts in a reproducible manner at the average population level for five replicated cultures with respect to changes in heterologous insulin production and intracellular proteome under prolonged glucose-limited conditions (Wright et al., 2020). Diversity of adaptation varies as a function of the distribution of fitness effects between beneficial outcomes (Gresham et al., 2008). Thus, if an adaptive path confers a much greater selective advantage compared to other selective outcomes, that path will be highly reproducible. Due to the reproducibility of the phenotypic outcome of chemostat cultivations with the *initial cell clone,* we expected the adaptation to be driven by the selection of a single clone with a large relative selective advantage. In the present study, however, we demonstrate that the adaptation is not caused by the selection towards a single beneficial phenotype. Instead, the isogenic strain differentiated into subpopulations and reproducibly established three main subpopulations after 271 hours of glucose-limited growth (Figure 3A). Our results indicate that three apparent phenotypes underlie the adaptive response observed at the bulk level (Figure 7). We speculate that this phenotypic heterogeneity is a result of different mechanisms to increase fitness. *Population 1* has increased fitness by downregulating heterologous insulin production most likely by decreasing the plasmid copy number whereas *Population 2* has adapted towards the glucose-limited conditions. *Population 3* seems to be a response to both the burden of the heterologous insulin production and the glucose-limited conditions having a phenotype with reduced insulin productivity and pseudo-hyphal growth.

**Figure 7:**
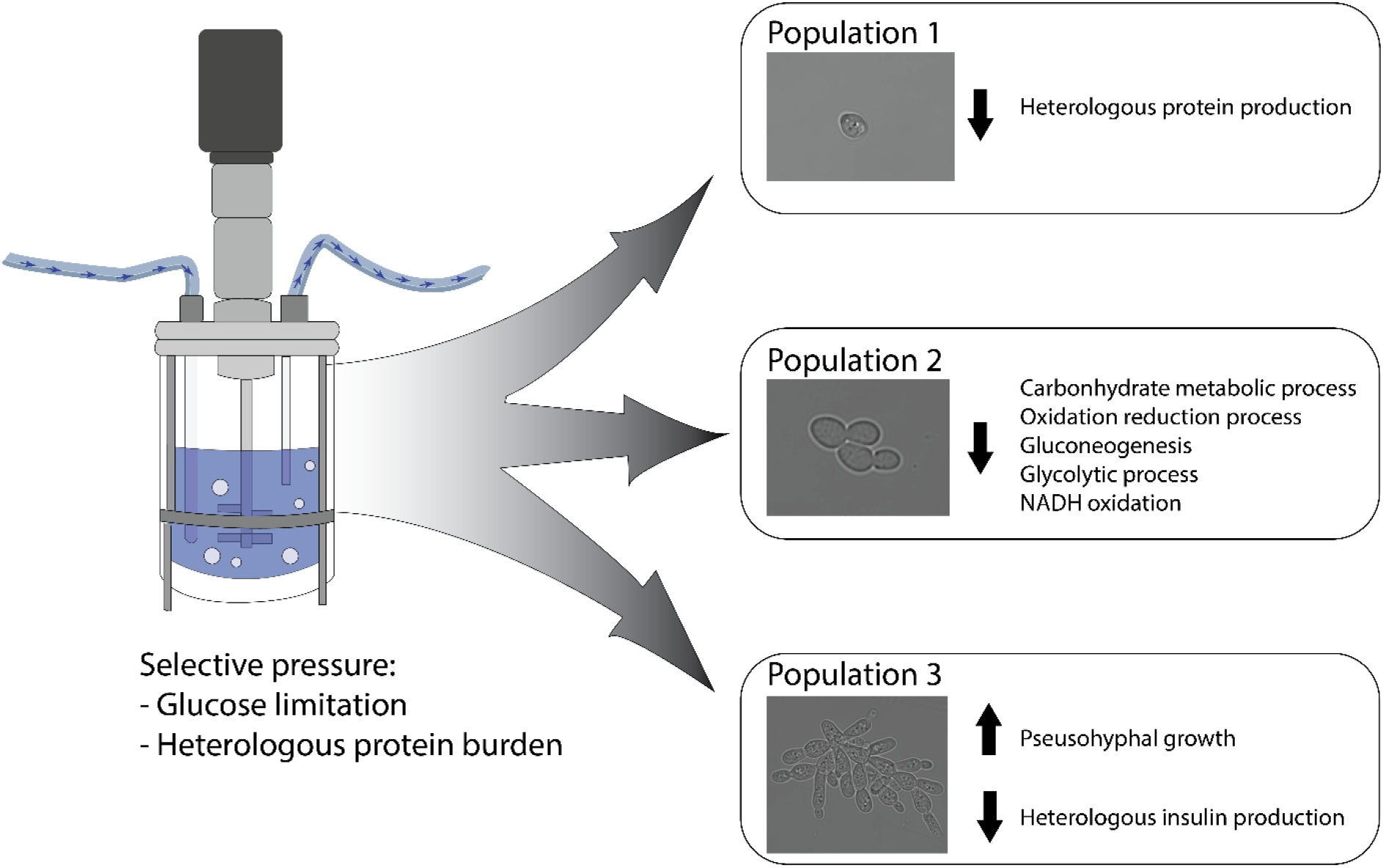
The adaptive outcome observed at the culture level is a mixed response of at least three apparent phenotypes. We speculate that Population 1 is mainly an outcome of the selective pressure of the glucose-limited conditions whereas Population 2 is a reaction towards the protein burden of the heterologous insulin production. Population 3 seems to be a product of both the selective pressure of the heterologous insulin burden and the glucose-limited conditions.

Adaptation and evolution in chemostats have been highly investigated at the average population level. However, only few studies investigate adaptation in terms of isogenic cells differentiating into phenotypic subpopulations (Kundu et al., 2020). Our results highlight the importance of considering population heterogeneity when studying adaptation as bulk adaptive outcomes observed at the culture level can be a mixed response composed of different phenotypes or subpopulations.

## 4. Conclusion

In this study, we investigated adaptation at the single-cell level and showed that an isogenic, recombinant *S. cerevisiae* strain differentiated into three subpopulations: (1) cells that drastically reduced insulin production (23 %), (2) cells with reduced enzymatic capacity of the central carbon metabolism (46 %), (3) cells that exhibited pseudohyphal growth (31 %). This indicates that the bulk adaptive outcome is a mixed response composed of at least three apparent subpopulations emerged as a response to the selective pressure of the chemostat and the burden of the heterologous insulin production.

## 5. Acknowledgements

This work received funding from Innovation Fund Denmark, case no. 7038-00165B, and the Novo Nordisk R&D STAR Fellowship Programme.

The authors Naia Risager Wright and Nanna Petersen Rønnest were employed by the company Novo Nordisk A/S. The remaining authors declare that the research was conducted in the absence of any commercial or financial relationships that could be construed as a potential conflict of interest.

