## Supplementary materials for "Emergence of phenotypically distinct subpopulations is a factor in adaptation of recombinant *Saccharomyces cerevisiae* under glucose-limited conditions"

### 1. Forward scatter area

The forward scatter area (FSC-A) was used as a metric for separation of subpopulations with respect to morphology. Over time the *initial cells clone* differentiates into three main morphological phenotypes (Figure S1).

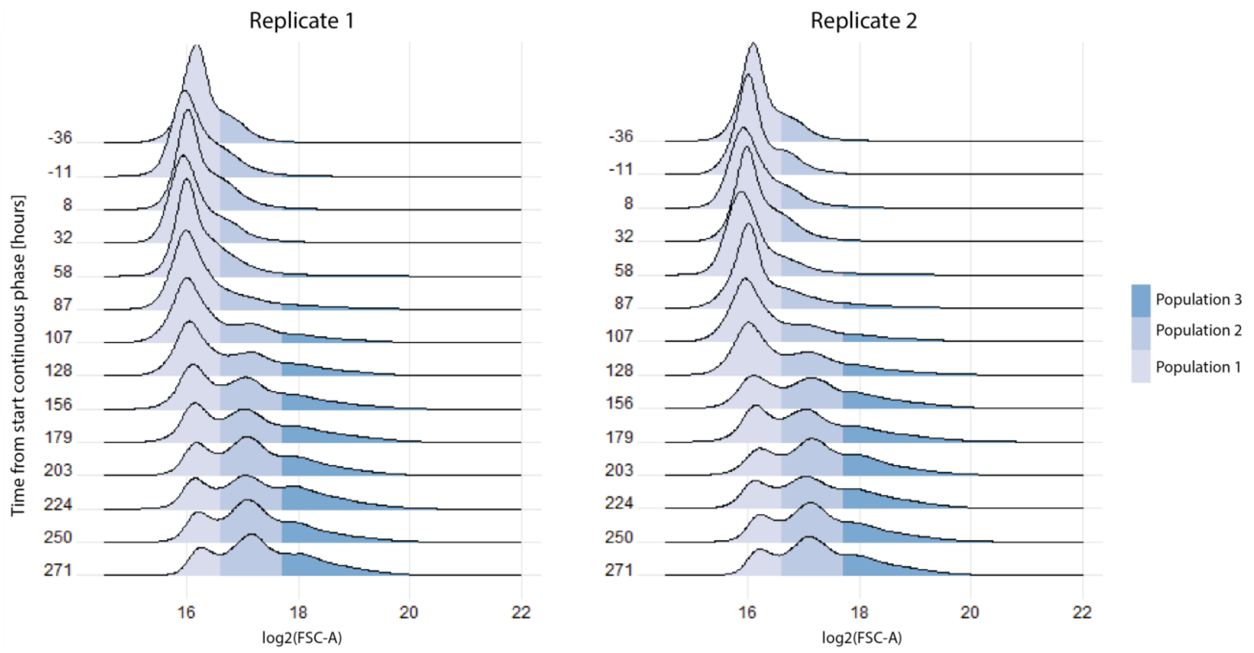


*Figure S1: Density plots of log2(FSC-A) for two cultivations with the initial cell clone at different time points in the cultivation. The cells are color coded with respect to FSC-A: Population 1 (log2(*FSC-A) <16.6), *Population 2 (16.6< log2(*FSC-A) <17.7*)* and *Population 3 (log2(FSC-A) > 17.7). 100,000 cells were analyzed in each sample.*

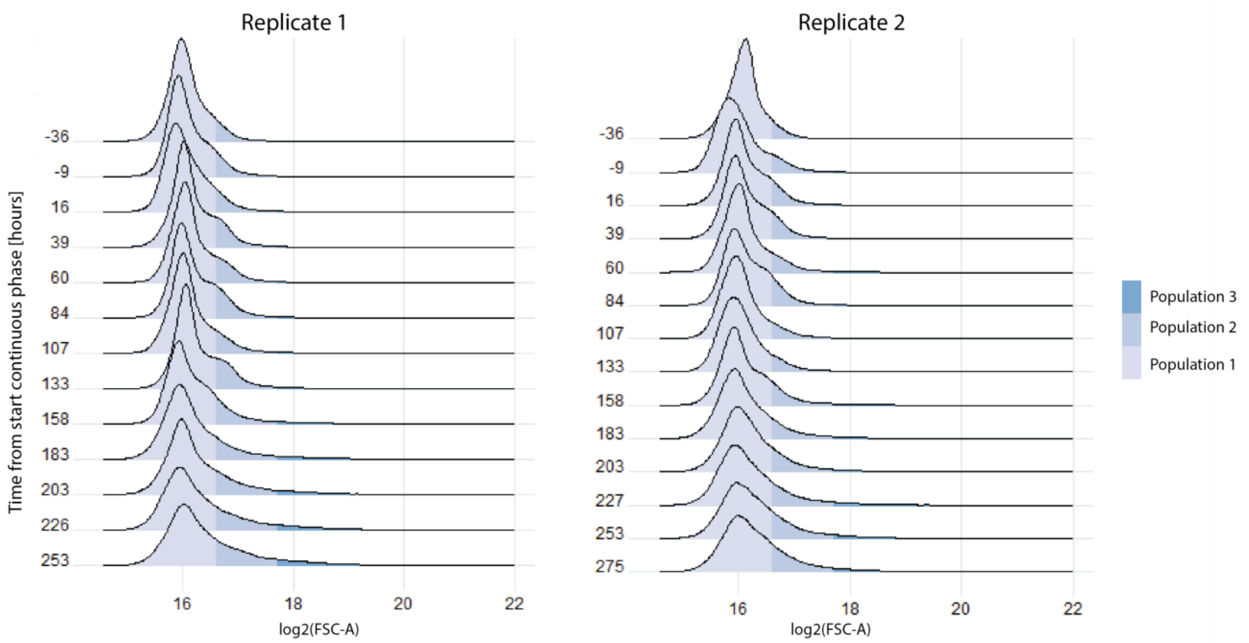


*Figure S2: Density plots of log2(FSC-A) for two cultivations with Population 1 at different time points in the cultivation. The cells are color coded with respect to FSC-A: Population 1 (log2(*FSC-A) <16.6), *Population 2 (16.6< log2(*FSC-A) <17.7*)* and *Population 3 (log2(FSC-A) > 17.7). 100,000 cells were analyzed in each sample.*

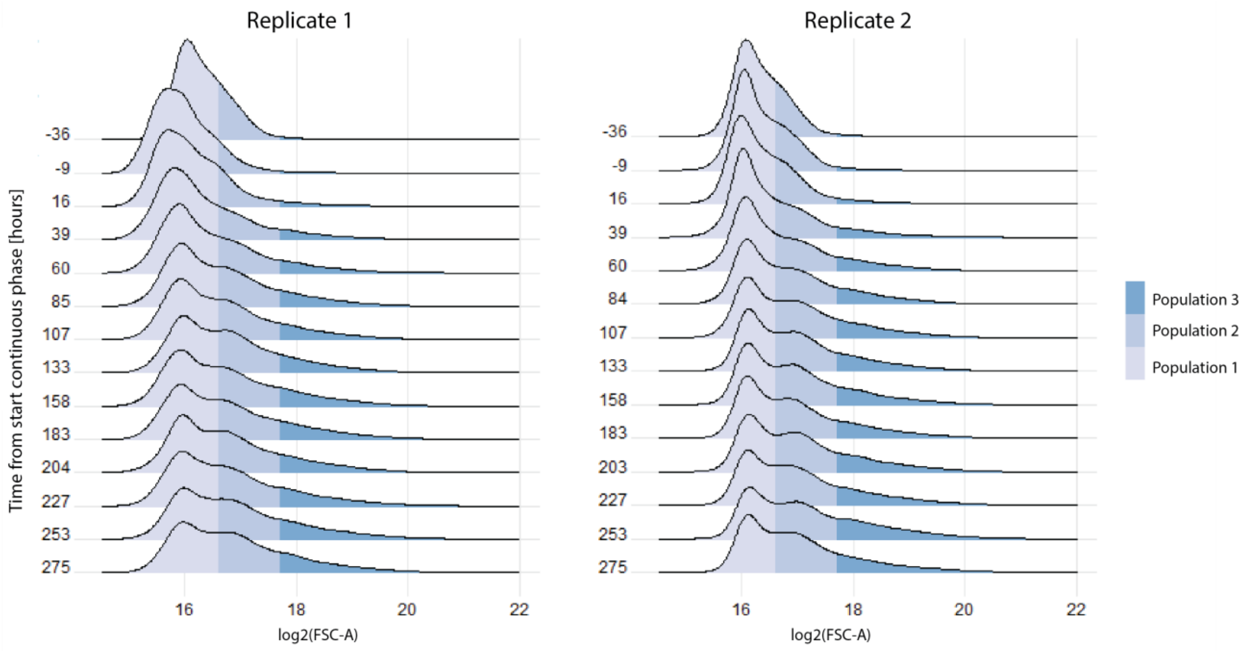


*Figure S3: Density plots of log2(FSC-A) for two cultivations with Population 2 at different time points in the cultivation. The cells are color coded with respect to FSC-A: Population 1 (log2(*FSC-A) <16.6), *Population 2 (16.6< log2(*FSC-A) <17.7) and *Population 3 (log2(FSC-A) > 17.7). 100,000 cells were analyzed in each sample.*

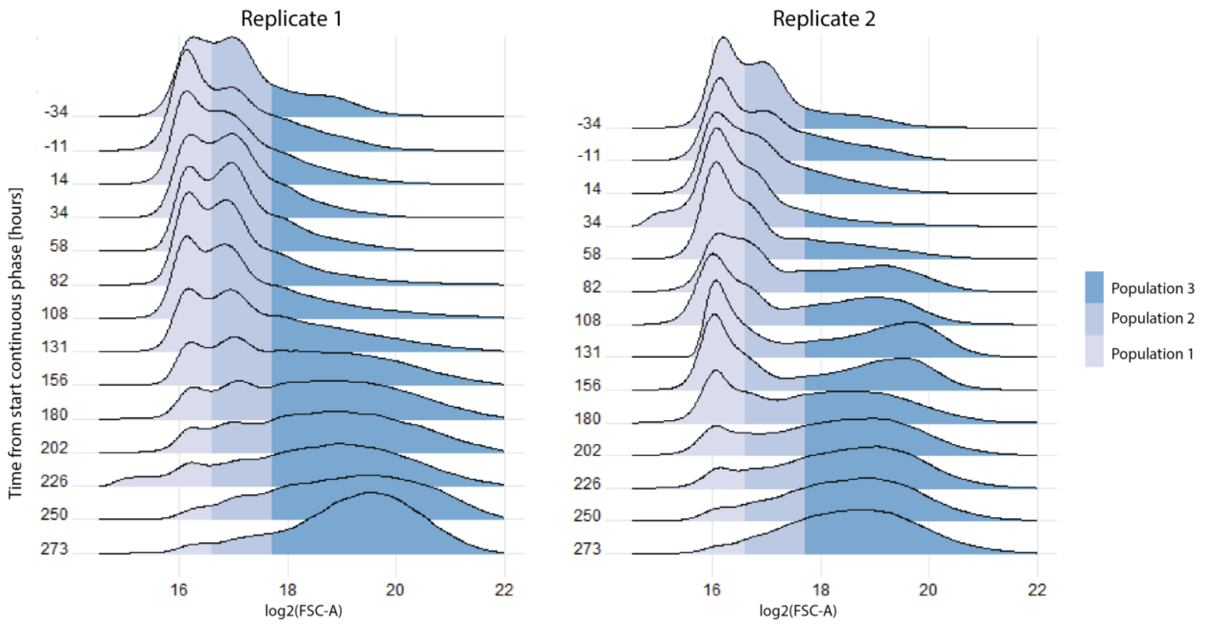


*Figure S4: Density plots of log2(FSC-A) for two cultivations with Population 3 at different time points in the cultivation. The cells are color coded with respect to FSC-A: Population 1 (log2(*FSC-A) <16.6), *Population 2 (16.6< log2(*FSC-A) <17.7) and *Population 3 (log2(FSC-A) > 17.7). 100,000 cells were analyzed in each sample.*

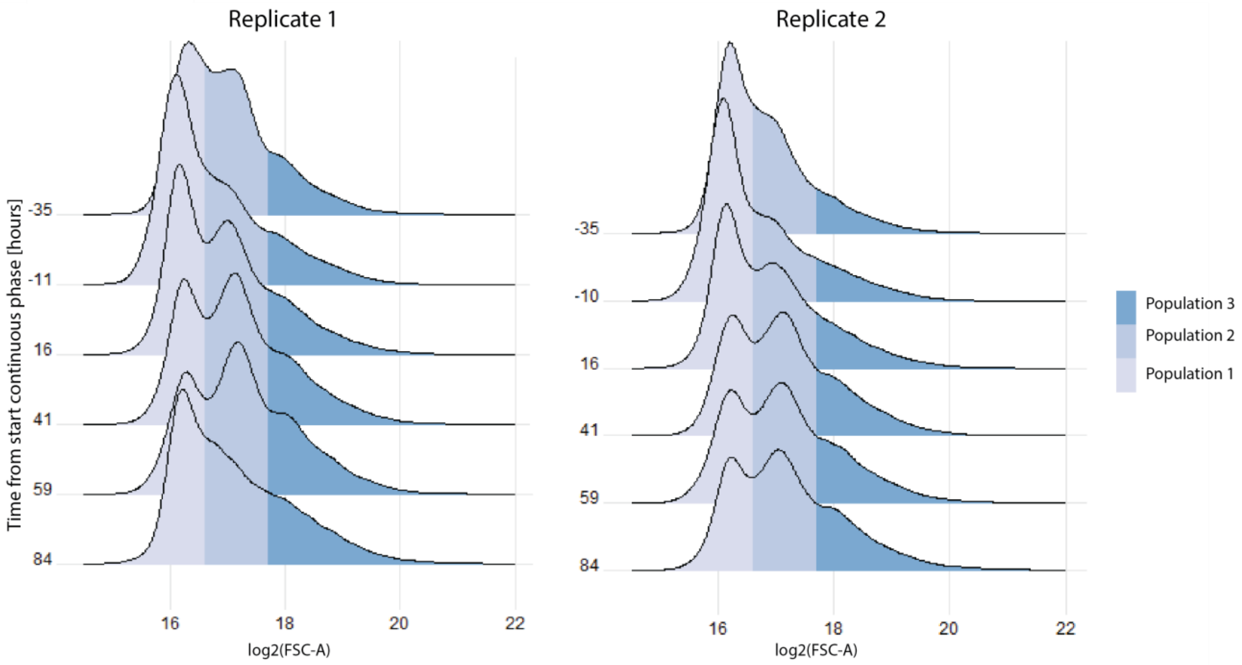


*Figure S5: Density plots of log2(FSC-A) for two cultivations with the end sample cells at different time points in the cultivation. The cells are color coded with respect to FSC-A: Population 1 (log2(*FSC-A) <16.6)*, Population 2 (16.6< log2(*FSC-A) <17.7) and *Population 3 (log2(FSC-A) > 17.7). 100,000 cells were analyzed in each sample.*

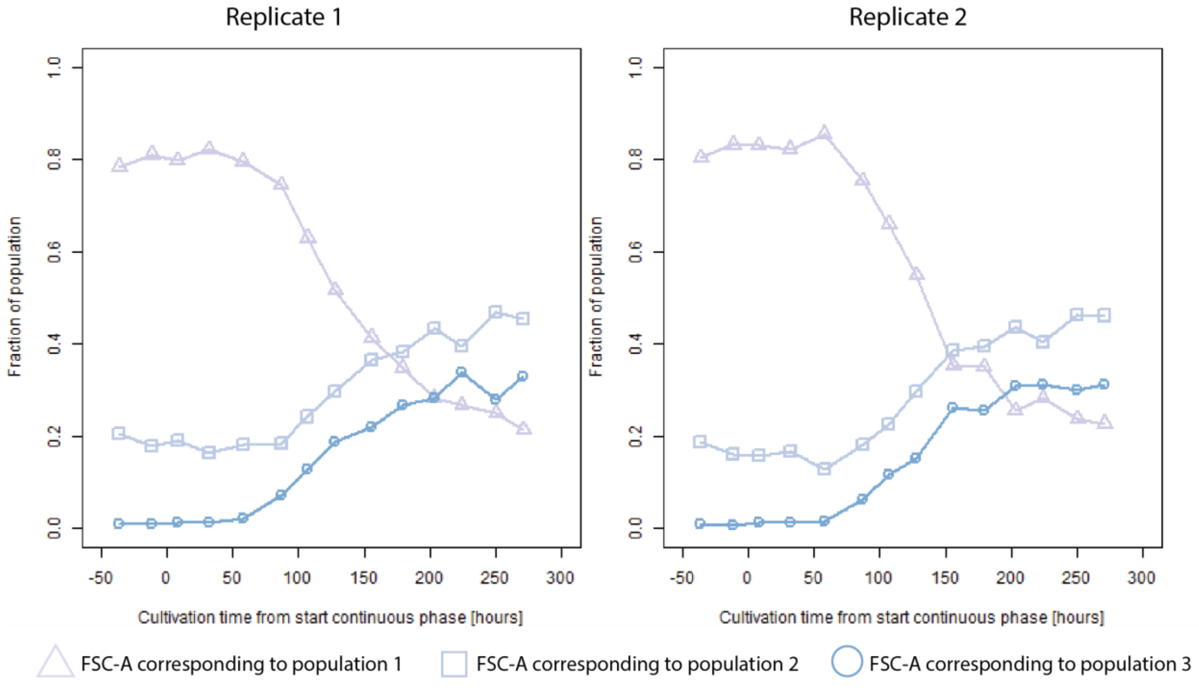


*Figure S6: Fraction of cells with FSC-A corresponding to Population 1 (log2(*FSC-A) <16.6)*, Population 2 (16.6< log2(*FSC-A) <17.7) or *Population 3 (log2(FSC-A) > 17.7) for two cultivations with the initial cell clone at different time points in the cultivation.*

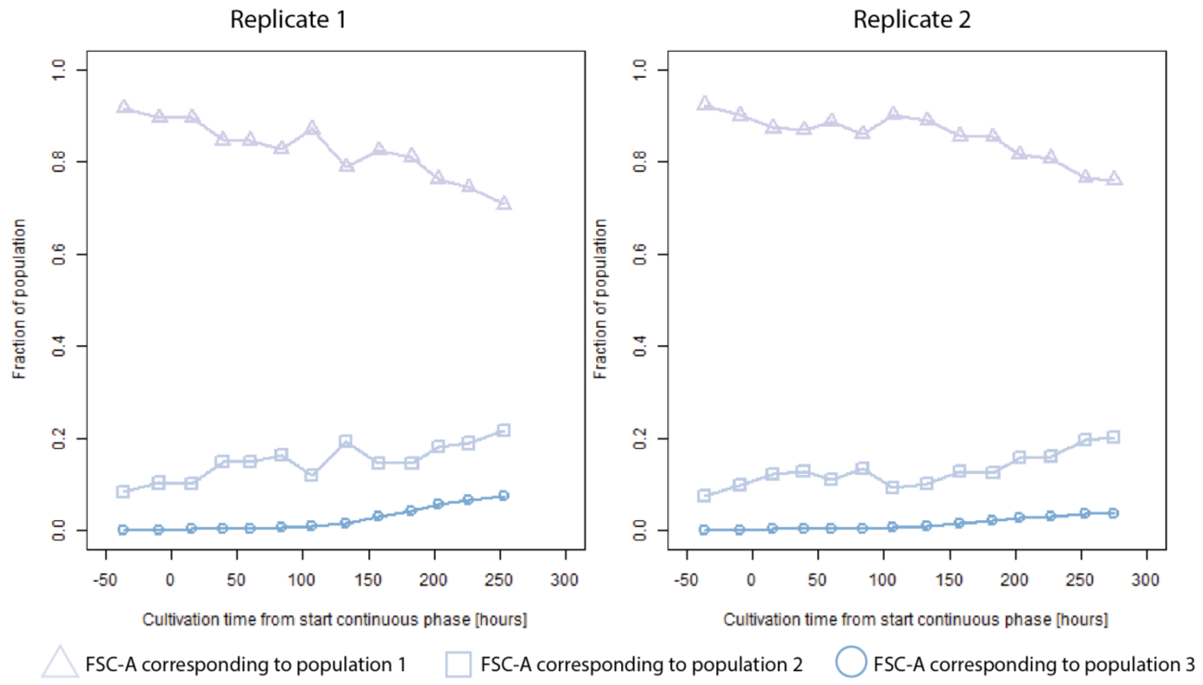


*Figure S7: Fraction of cells with FSC-A corresponding to Population 1 (log2(*FSC-A) <16.6), *Population 2 (16.6< log2(*FSC-A) <17.7) or *Population 3 (log2(FSC-A) > 17.7) for two cultivations with Population 1 at different time points in the cultivation.*

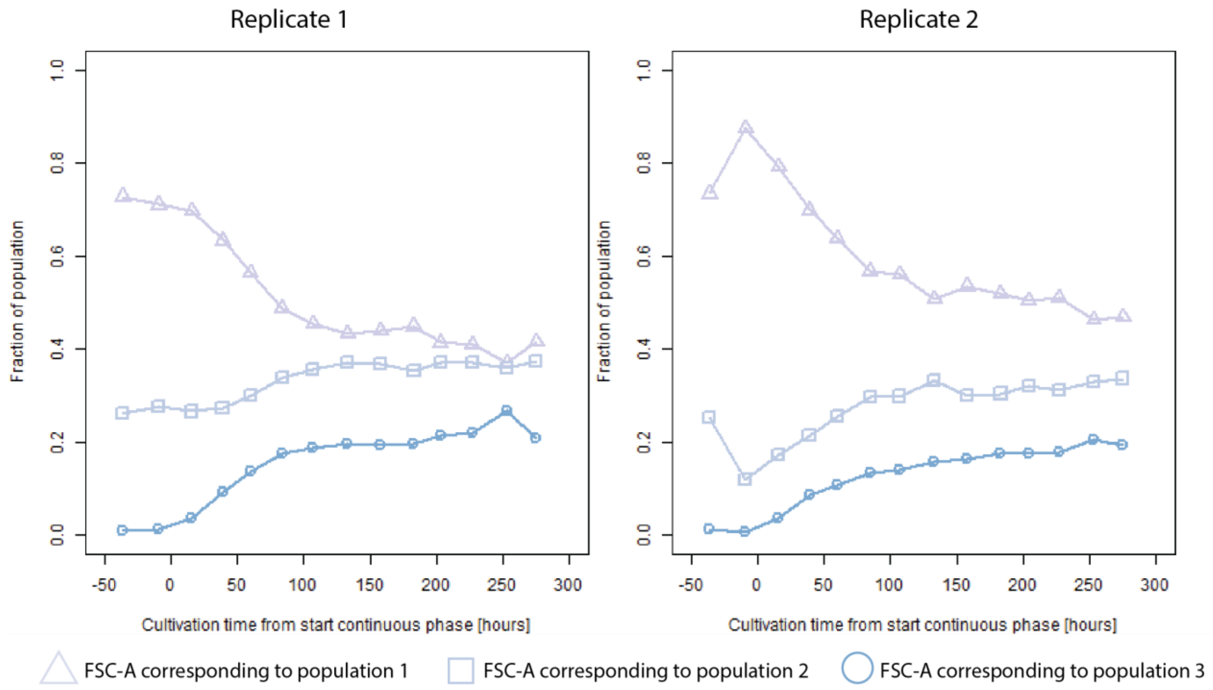


*Figure S8: Fraction of cells with FSC-A corresponding to Population 1 (log2(*FSC-A) <16.6), *Population 2 (16.6< log2(*FSC-A) <17.7) or *Population 3 (log2(FSC-A) > 17.7) for two cultivations with Population 2 at different time points in the cultivation.*

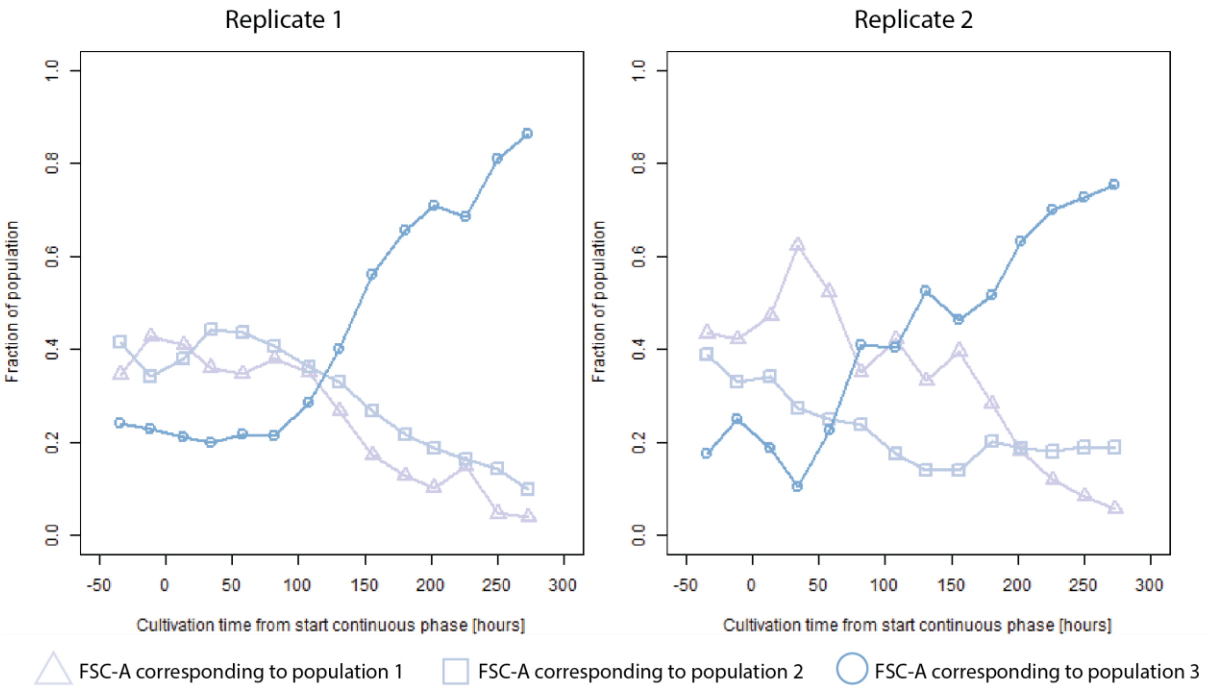


*Figure S9: Fraction of cells with FSC-A corresponding to Population 1 (log2(*FSC-A) <16.6), *Population 2 (16.6< log2(*FSC-A) <17.7) or *Population 3 (log2(FSC-A) > 17.7) for two cultivations with Population 3 at different time points in the cultivation.*

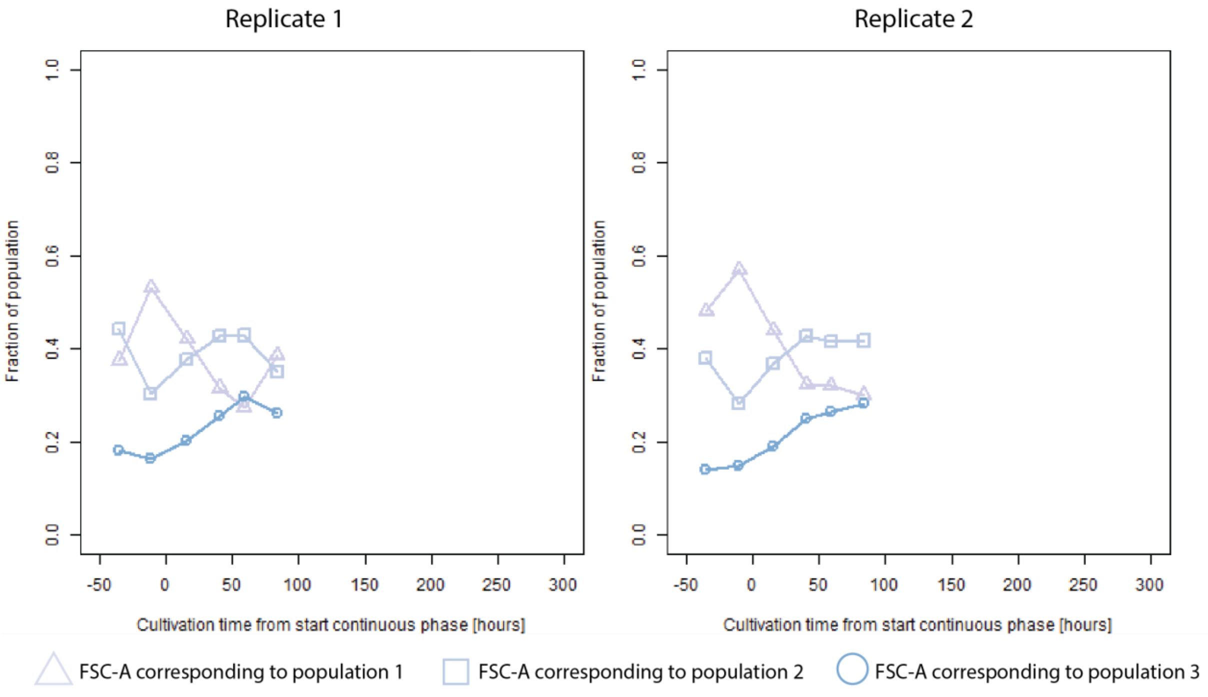


*Figure S10: Fraction of cells with FSC-A corresponding to Population 1 (log2(*FSC-A) <16.6), *Population 2 (16.6< log2(*FSC-A) <17.7) or *Population 3 (log2(FSC-A) > 17.7) for two cultivations with the end sample cells at different time points in the cultivation.*

#### 2. Cultivations with pulse feeding

To investigate whether the emergence of the subpopulations was a reaction to the glucose-limited conditions, we performed cultivations continuously exposed to glucose pulses. The setup of the cultivations were the same as described in the Materials and Methods section but instead of a continuous media supply, the media is dosed in pulses. Thus, the media, which should be added to the reactor in a time interval of 84 seconds in order to obtain a dilution rate of 0.1 $h^{-1}$ are instead added in 22 seconds followed by a 62 seconds pause with no media addition. No subpopulations were observed in these cultivations (Figure S11,S12). No decline in the bulk measured insulin production was observed in these cultivations (Figure S13). This correspond to our previous observations (Wright et al. 2020).

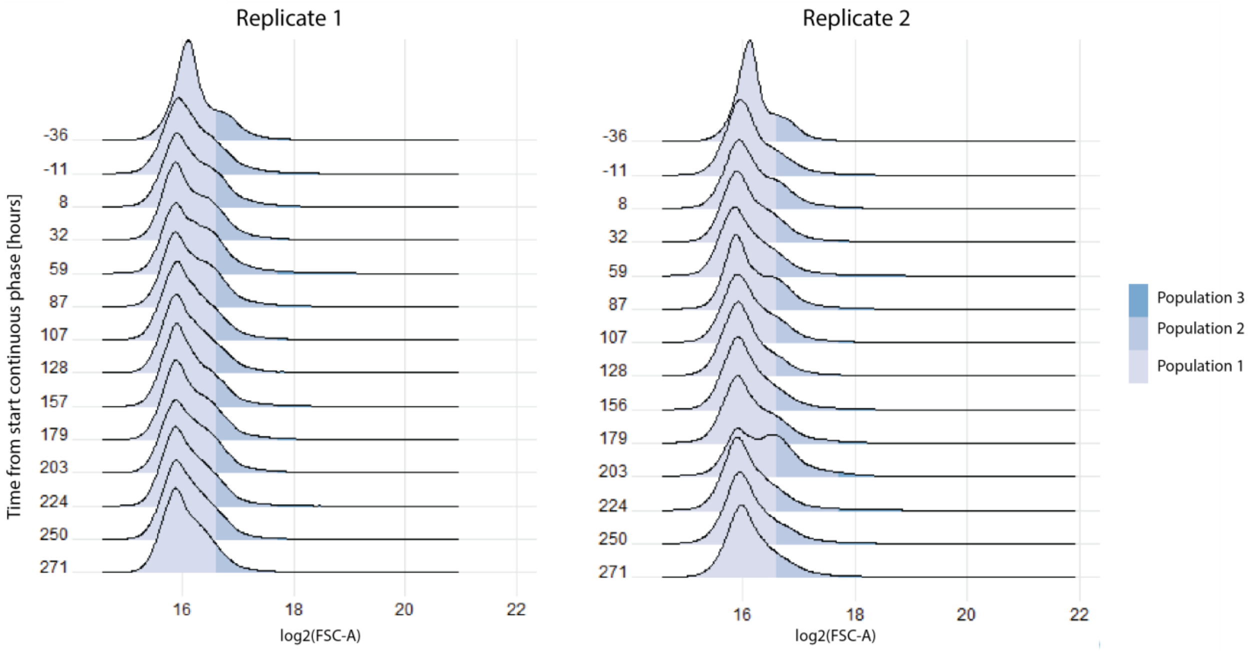


*Figure S11 Density plots of log2(FSC-A) for two cultivations with Population1 at different time points in the cultivation. Media was supplied in pulses of 22 seconds followed by and pause of 62 seconds with no media addition. The cells are color coded with respect to FSC-A: Population 1 (log2(*FSC-A) <16.6)*, Population 2 (16.6< log2(*FSC-A) <17.7*)* and *Population 3 (log2(FSC-A) > 17.7). 100,000 cells were analyzed in each sample.*

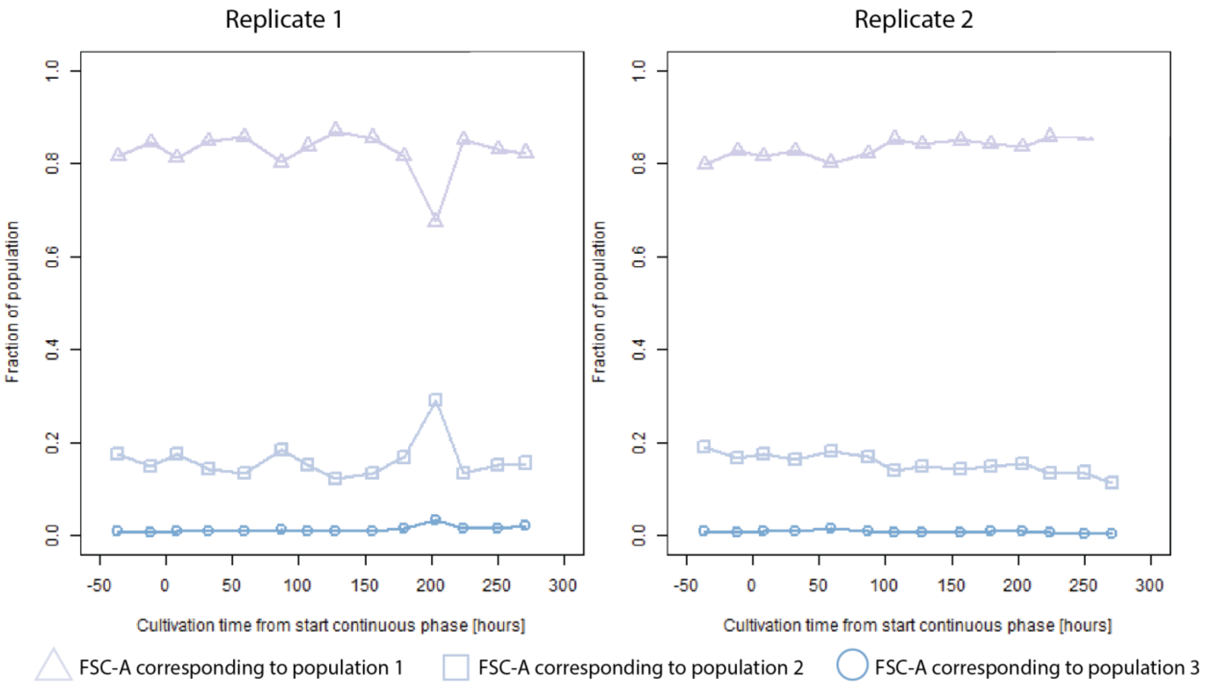


*Figure S12: Fraction of cells with FSC-A corresponding to Population 1 (log2(*FSC-A) <16.6)*, Population 2 (16.6<log2(* FSC-A) <17.7) or *Population 3 (log2(FSC-A) > 17.7) for two cultivations with the initial cell clone at different time points in the cultivation. Media is supplied in pulses of 22 seconds followed by and pause of 62 seconds with no media addition.*

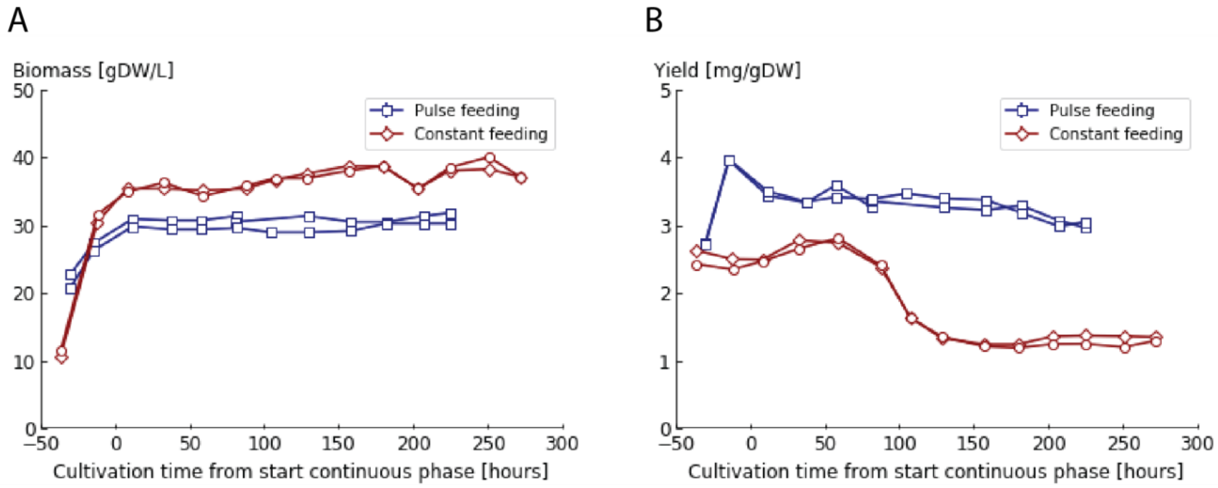


*Figure S13: Biomass concentration (A) and insulin yield (B) as function of cultivation time for the cultivations with the initial cell clone with pulse feeding and constant feeding respectively.*

#### 3. Variance in reinitiated chemostats

*Population 3* was defined as cells with a log2(FSC-A) larger than 17.5. Thus, this subpopulation spanned over a larger range of cells with respect to FSC-A compared to *Population 1* and *Population 2* (Figure 3A). This may explain the larger variation observed between replicates with respect to intracellular protein levels and insulin production compared to *Population 1* and *Population 2* (Figure S14).

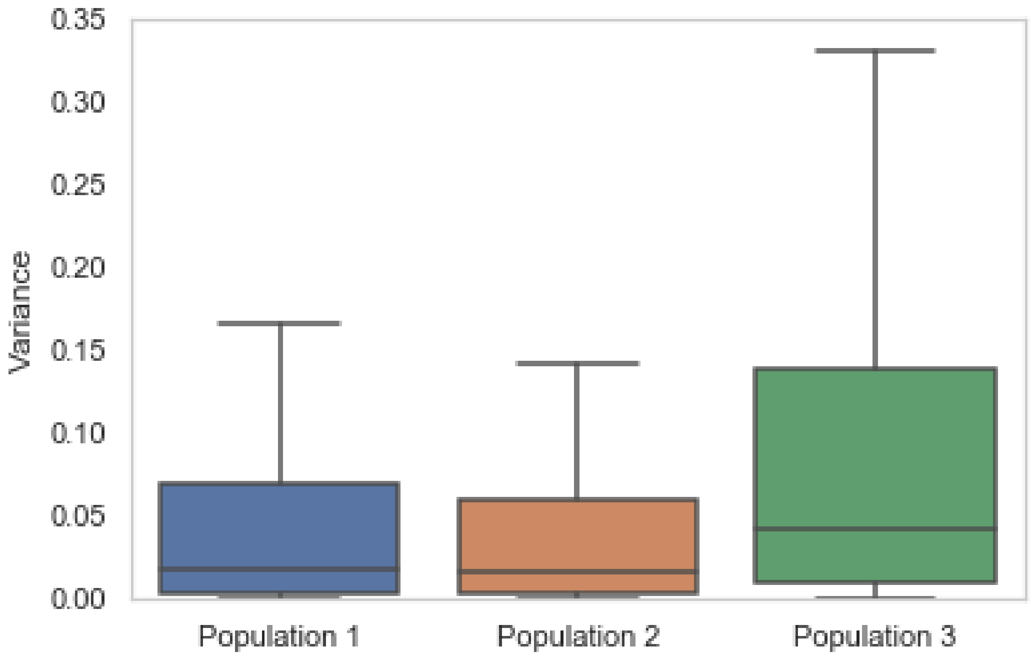


*Figure S14:* Boxplot of variances for all proteins measured after 48 hours of chemostat growth between two replicated cultivations of *Population 1*, *Population 2* and *Population 3* respectively.

### 4. Intracellular proteome

*Table S1: Overview of analyzed proteomics samples from reinitiated chemostat cultivations with the FACS sorted populations.*

| Sample name | Fermentation batch | Strain | Time from start continuous phase [hours] | Proteomics batch |
|---|---|---|---|---|
| DCL209_P1 | DCL209 | Population 1 | 48 | Proteomics_Population_1_2 |
| DCL209_P2 | DCL209 | Population 1 | 91 | Proteomics_Population_1_2 |
| DCL209_P3 | DCL209 | Population 1 | 137 | Proteomics_Population_1_2 |
| DCL209_P4 | DCL209 | Population 1 | 254 | Proteomics_Population_1_2 |
| DCL209_P5 | DCL209 | Population 1 | 276 | Proteomics_Population_1_2 |
| DCL210_P1 | DCL210 | Population 1 | 48 | Proteomics_Population_1_2 |
| DCL210_P2 | DCL210 | Population 1 | 91 | Proteomics_Population_1_2 |
| DCL210_P3 | DCL210 | Population 1 | 137 | Proteomics_Population_1_2 |
| DCL210_P4 | DCL210 | Population 1 | 254 | Proteomics_Population_1_2 |
| DCL210_P5 | DCL210 | Population 1 | 276 | Proteomics_Population_1_2 |
| DCL211_P1 | DCL211 | Population 2 | 48 | Proteomics_Population_1_2 |
| DCL211_P2 | DCL211 | Population 2 | 91 | Proteomics_Population_1_2 |
| DCL211_P3 | DCL211 | Population 2 | 137 | Proteomics_Population_1_2 |
| DCL211_P4 | DCL211 | Population 2 | 254 | Proteomics_Population_1_2 |
| DCL211_P5 | DCL211 | Population 2 | 276 | Proteomics_Population_1_2 |
| DCL212_P1 | DCL212 | Population 2 | 48 | Proteomics_Population_1_2 |
| DCL212_P2 | DCL212 | Population 2 | 91 | Proteomics_Population_1_2 |
| DCL212_P3 | DCL212 | Population 2 | 137 | Proteomics_Population_1_2 |
| DCL212_P4 | DCL212 | Population 2 | 254 | Proteomics_Population_1_2 |
| DCL212_P5 | DCL212 | Population 2 | 276 | Proteomics_Population_1_2 |
| DCL221_P1 | DCL221 | Population 3 | 37 | Proteomics_Population_3 |
| DCL221_P2 | DCL221 | Population 3 | 203 | Proteomics_Population_3 |
| DCL221_P3 | DCL221 | Population 3 | 254 | Proteomics_Population_3 |
| DCL221_P4 | DCL221 | Population 3 | 131 | Proteomics_Population_3 |
| DCL222_P1 | DCL221 | Population 3 | 37 | Proteomics_Population_3 |
| DCL222_P2 | DCL221 | Population 3 | 203 | Proteomics_Population_3 |
| DCL222_P3 | DCL221 | Population 3 | 254 | Proteomics_Population_3 |
| DCL222_P4 | DCL221 | Population 3 | 131 | Proteomics_Population_3 |

*Table S2: Overview of proteomics samples from the cultivation with the Initial cell clone used for comparison with the proteome of Population 1-3 after 48 hours of chemostat culture. Data were obtained from* (Wright et al. 2020).

| Sample no | Cultivation equipment | Cultivation time from start continuous phase [hours] | Strain ID | Cultivation batch no. _sample_no |
|---|---|---|---|---|
| Abundances (Normalized): F19 | Single compartment system | 31.42 | C.U17 | DCL136_P1 |
| Abundances (Normalized): F25 | Single compartment system | 8.84 | C.U17 | DCL137_P1 |
| Abundances (Normalized): F56 | Single compartment system | 33.92 | C.U17 | DCL141_P2 |
| Abundances (Normalized): F63 | Single compartment system | 34.00 | C.U17 | DCL142_P2 |

*Table S3: Differentially expressed proteins between population 1 and population 2 in the end of chemostat cultivations (254 hours of chemostat growth), (log2 fold change > 0.5 or log2 fold change < -0.5, q-value <0.05).*

| Protein | q-value |
|---|---|
| TDH2 | 0.039 |
| TDH1 | 0.021 |
| GPH1 | 0.026 |
| YDL124W | 0.048 |
| URA3 | 0.026 |
| GCY1 | 0.021 |
| TRR1 | 0.026 |
| GND2 | 0.021 |
| ALT1 | 0.026 |
| RNR4 | 0.039 |
| HIS3 | 0.009 |
| Insulin | 0.021 |
| PBS2 | 0.026 |
| EGT2 | 0.026 |
| MF(ALPHA)1 | 0.021 |
| FAT3; YKL187C | 0.026 |

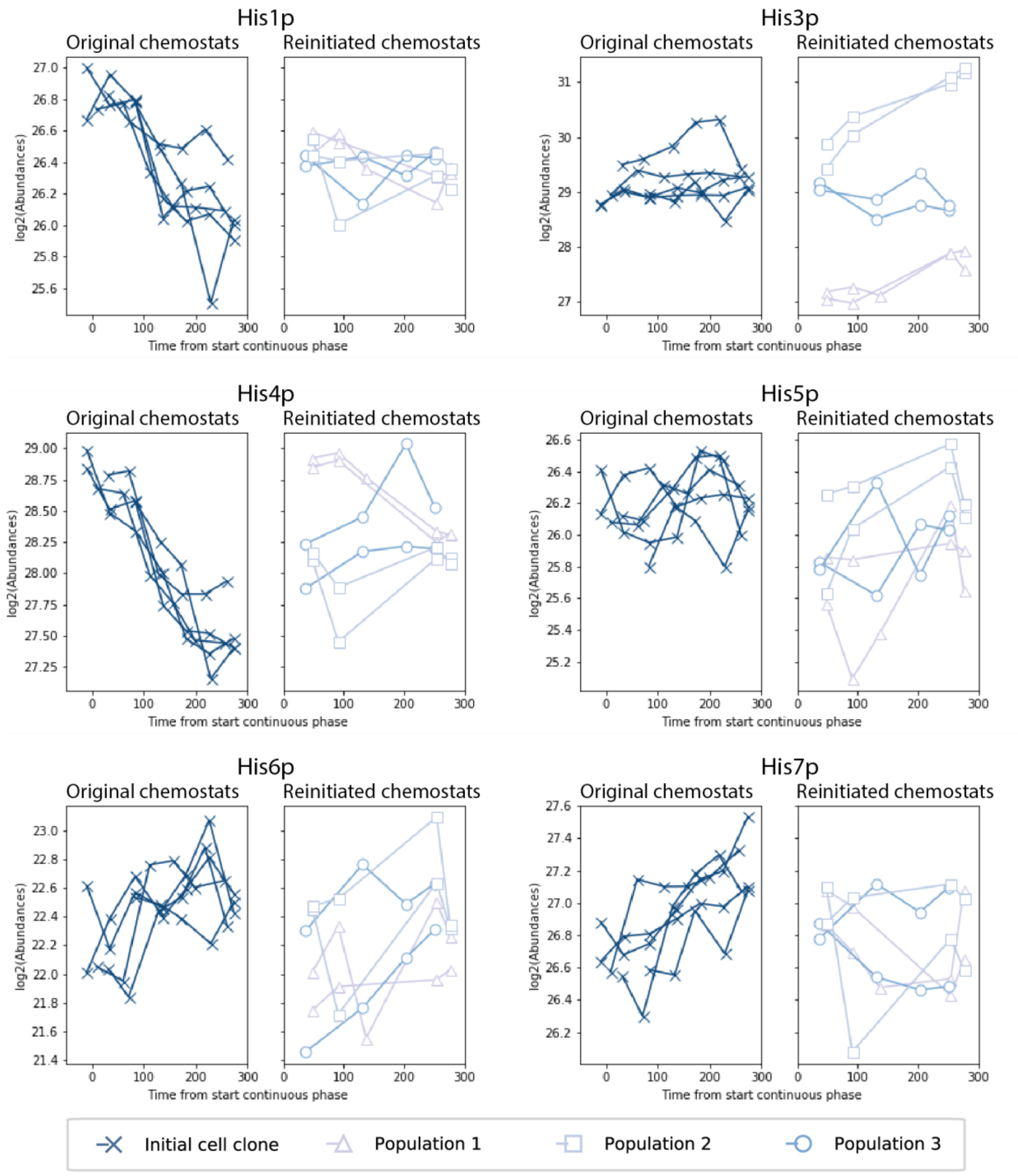


*Figure S15: Measured enzymes involved in the histidine biosynthetic pathway.*

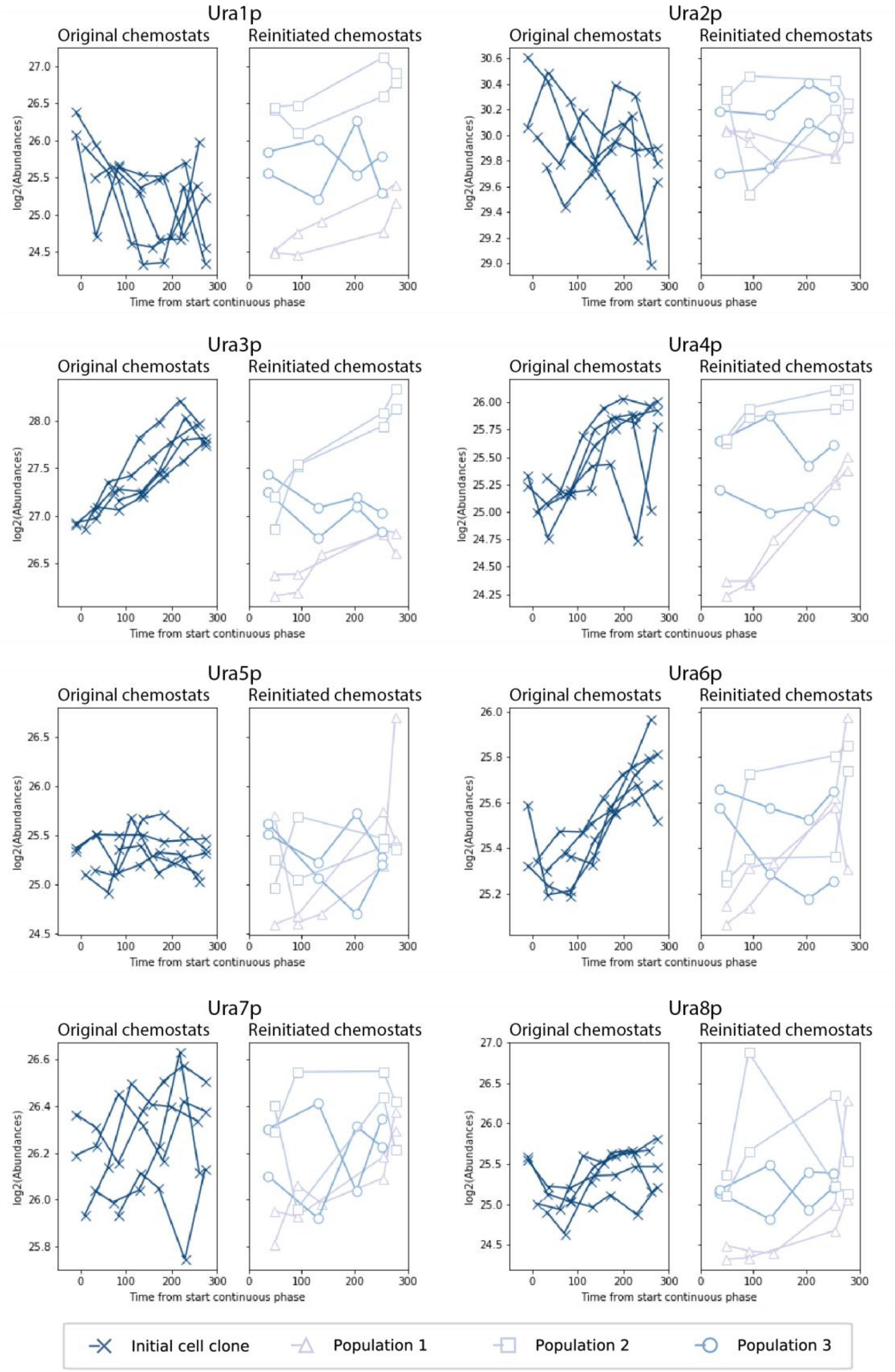


*Figure S16: Measured enzymes involved in the uracil biosynthetic pathway.*

#### 5. Biomass concentration

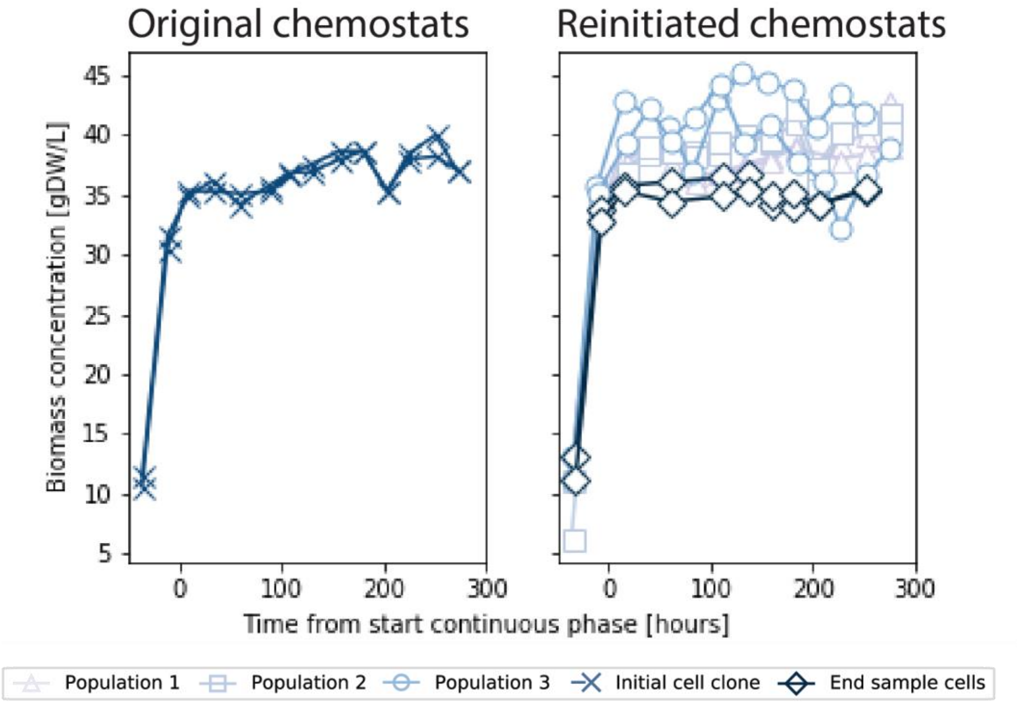


*Figure S17: Biomass concentration as function of cultivation time for chemostat cultivations with the initial cell clone, Population 1, Population 2, Population 3 and the End sample cells.*
